# Automated Biomarker Candidate Discovery in Imaging Mass Spectrometry Data Through Spatially Localized Shapley Additive Explanations

**DOI:** 10.1101/2020.12.23.424201

**Authors:** Leonoor E.M. Tideman, Lukasz G. Migas, Katerina V. Djambazova, Nathan Heath Patterson, Richard M. Caprioli, Jeffrey M. Spraggins, Raf Van de Plas

## Abstract

The search for molecular species that are differentially expressed between biological states is an important step towards discovering promising biomarker candidates. In imaging mass spectrometry (IMS), performing this search manually is often impractical due to the large size and high-dimensionality of IMS datasets. Instead, we propose an interpretable machine learning workflow that automatically identifies biomarker candidates by their mass-to-charge ratios, and that quantitatively estimates their relevance to recognizing a given biological class using Shapley additive explanations (SHAP). The task of biomarker candidate discovery is translated into a feature ranking problem: given a classification model that assigns pixels to different biological classes on the basis of their mass spectra, the molecular species that the model uses as features are ranked in descending order of relative predictive importance such that the top-ranking features have a higher likelihood of being useful biomarkers. Besides providing the user with an experiment-wide measure of a molecular species’ biomarker potential, our workflow delivers spatially localized explanations of the classification model’s decision-making process in the form of a novel representation called SHAP maps. SHAP maps deliver insight into the spatial specificity of biomarker candidates by highlighting in which regions of the tissue sample each feature provides discriminative information and in which regions it does not. SHAP maps also enable one to determine whether the relationship between a biomarker candidate and a biological state of interest is correlative or anticorrelative. Our automated approach to estimating a molecular species’ potential for characterizing a user-provided biological class, combined with the untargeted and multiplexed nature of IMS, allows for the rapid screening of thousands of molecular species and the obtention of a broader biomarker candidate shortlist than would be possible through targeted manual assessment. Our biomarker candidate discovery workflow is demonstrated on mouse-pup and rat kidney case studies.

**Highlights:**

- Our workflow automates the discovery of biomarker candidates in imaging mass spectrometry data by using state-of-the-art machine learning methodology to produce a shortlist of molecular species that are differentially expressed with regards to a user-provided biological class.
- A model interpretability method called Shapley additive explanations (SHAP), with observational Shapley values, enables us to quantify the local and global predictive importance of molecular species with respect to recognizing a user-provided biological class.
- By providing spatially localized explanations for a classification model’s decision-making process, SHAP maps deliver insight into the spatial specificity of biomarker candidates and enable one to determine whether (and where) the relationship between a biomarker candidate and the class of interest is correlative or anticorrelative.

## l Introduction

A biomarker can generally be considered an objectively measurable indicator of a specific biological state or disease condition [1, 2]. Biomarkers can be used to differentiate between anatomical structures, cell types, and disease states, and lend themselves to the screening, diagnosis, and monitoring of disease, the identification of new drug targets, and the assessment of therapeutic response [1, 3, 4]. In our work, the term “biomarker candidate” refers to a putative molecular biomarker (i.e. a chemical species) that is differentially expressed between biological states [2]. One technology for discovering such molecular markers at scale is mass spectrometry, which characterizes molecular species in terms of their mass-to-charge ratio (*m/z*). It was demonstrated in 2003 that matrix assisted laser desorption/ionization (MALDI) mass spectrometry could facilitate the discovery of diagnostic and prognostic biomarkers for lung cancer when applied directly to clinical patient samples [5, 6]. Imaging mass spectrometry (IMS) is a multiplexed, label-free imaging technology that uses mass spectrometry for the molecular mapping of tissues down to cellular resolution [7, 8, 9]. An IMS experiment involves collecting spatially localized mass spectra for each pixel in a grid of measurement locations across a sample surface [10, 11]. Each pixel has an associated mass spectrum and each mass spectrum plots the measured signal intensity, which is indicative of relative abundance, versus the analytes’ *m/z* values. The spatial distribution and relative abundance of an analyte can be visualized as an ion image, which plots the signal intensity measured for that analyte across all pixels of the sample’s surface [12, 13]. IMS is an excellent tool for biomarker discovery for the following three reasons: it is able to concurrently detect hundreds to thousands of analytes within a single experiment in an untargeted manner, it can probe analytes from a wide range of molecular classes (e.g. peptides, proteins, lipids, glycans, metabolites), and it enables the mapping of analytes’ spatial distributions in relation to the (patho)histology of tissue samples [14, 15]. There are several examples of IMS facilitating biomarker candidate discovery in cancer research. For example, MALDI IMS has been used to retrieve proteomic biomarker candidates for high-grade sarcomas [16] and melanomas [17]; and another IMS modality, called desorption electrospray ionization IMS, has been applied to the molecular study of brain cancer [18] and lung cancer [19], resulting in the discovery of diagnostic and prognostic biomarker candidates.

One way novel biomarker candidates can be discovered is by observing the differential expression of molecules between distinct sample classes (e.g. different cell types, different organs, different stages of a disease) [2, 20]. However, the large size and high-dimensionality of IMS datasets, which commonly yield several hundreds of thousands of pixels and several hundreds to thousands of molecular ions tracked per pixel, pose a challenge. Manually examining the spatial mapping of thousands of molecular species across the surface of a sample is laborious and risks introducing human subjectivity into the process, leading to results whose reproducibility cannot necessarily be guaranteed [12, 21]. The amount of data generated by IMS experiments is so large that it has become more efficient (and in many cases necessary) to computationally search for biomarker candidates among a multitude of ion intensity signals [22]. In this work, we suggest a machine learning (ML) workflow for performing biomarker candidate discovery that provides one with a shortlist of molecular species that are characteristic of the class for which biomarkers are sought. Our approach uses supervised ML models to classify mass spectra into different biological classes of interest and then uses state-of-the-art methods from the field of interpretable ML [23, 24, 25] to determine the discriminative relevance, and biomarker potential, of each molecular species.

In our work, an IMS dataset is represented by a data matrix *X* ∈ ℝ^*m*×*n*^ whose rows *x_i_* = *X*_(*i*,:)_, for *i* = 1, 2, 3…*m*, correspond to the mass spectra of the pixels making up the sample’s surface and whose columns *x ^j^* = *X*_(:,*j*)_, for *j* = 1, 2, 3…*n*, correspond to the *m/z* bins per spectrum. The *m* rows and *n* columns of *X* can be respectively referred to as observations and features. Classification is a form of supervised ML in which the observations *x_i_* are annotated with discrete class labels *y_i_* that represent user-provided knowledge related to these observations. Binary classification problems involve a positive class (e.g. diseased tissue), labeled as *y_i_* = +1, and a negative class (e.g. healthy tissue), labeled as *y_i_* = −1. The positive class is usually the class of interest: in our case studies, it is the class for which we want to discover biomarker candidates. Problems with multiple target classes (e.g. multiple cell types or functional tissue units) can be decomposed into multiple binary classification problems, each of which involve differentiating one class from the remaining classes. In the context of our work, classification is therefore the task of learning a multivariate function *f* ^⋆^ : ℝ^*n*^ → {−1, +1}, called a classification model, that assigns each pixel *x_i_* to a class according to the molecular information provided by its mass spectrum *x_i_*. Note a difference between the model’s class prediction 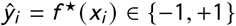 and the model’s raw output *f* (*x_i_*) ∈ ℝ, where *f* : ℝ^*n*^ → ℝ. The model’s prediction is the class label assigned to a particular observation *x_i_*, whereas the classification model’s raw output can be interpreted as the score (e.g. probability, log-odds ratio) of *x_i_* being assigned to the positive class. Figure 1a illustrates the process of building a classification model in IMS: a supervised ML algorithm fits a classification model to a labeled IMS dataset called the training dataset (i.e. mass spectra *x_i_* whose class membership *y_i_* is known). The resulting model can then be used to classify new unlabeled data (i.e. mass spectra *x_i_* whose class membership *y_i_* is unknown) as illustrated by Figure 1b. The performance of a classification model is measured by its ability to generalize, that is to correctly predict the labels for new data instances such that 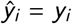.

**Figure 1.**
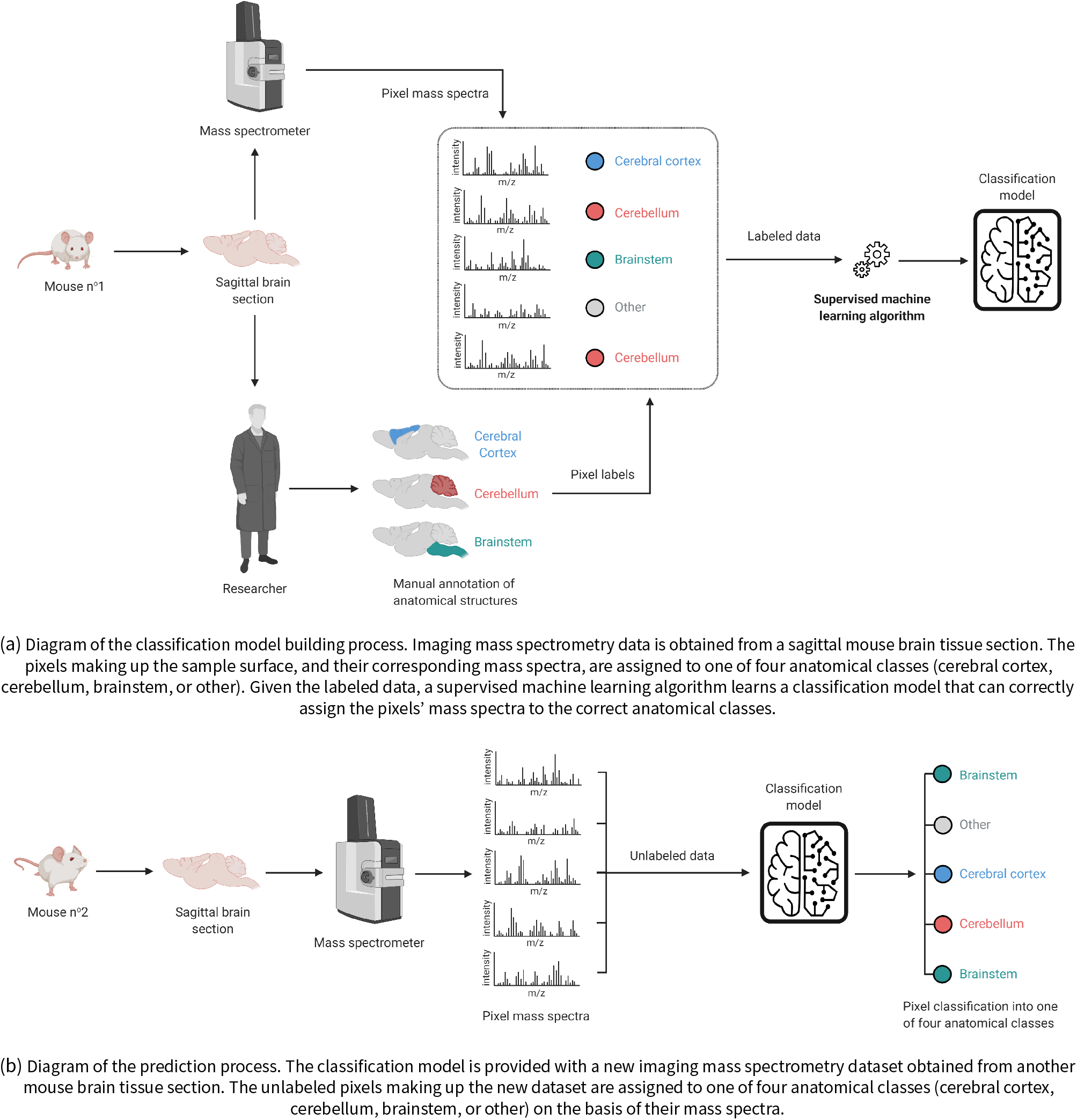
Diagrams of the classification model building and prediction processes in imaging mass spectrometry. Icons from [32, 33].

**Figure 2.**
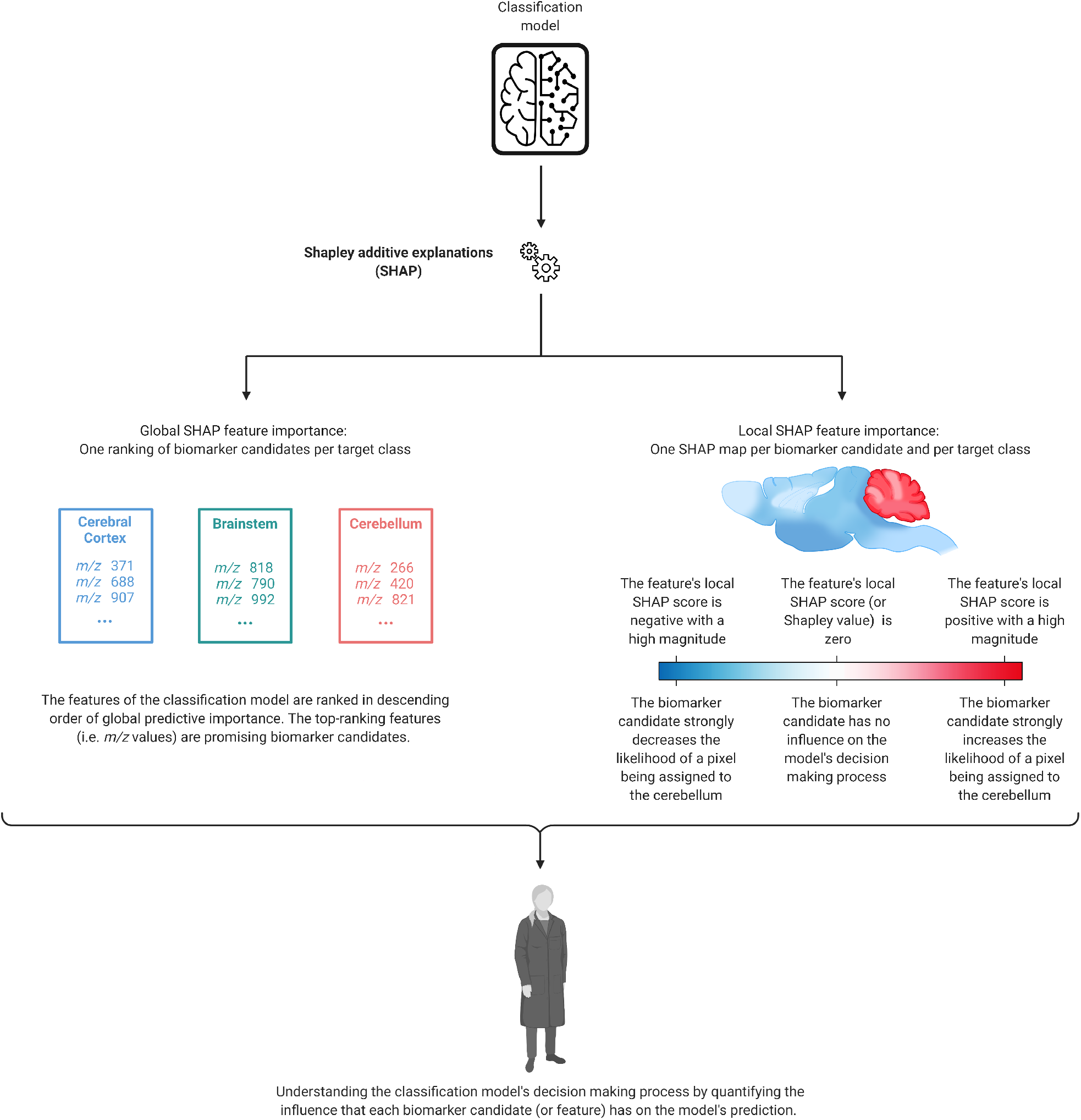
Diagram of the model interpretation process. SHAP is used to measure the local and global predictive importance of the features that the classification model from Figure 1 uses to assign the pixels making up the sample surface (and their corresponding mass spectra) to one of four different anatomical classes (cerebral cortex, cerebellum, brainstem, or other). The global SHAP scores provide an experiment-wide measure of each biomarker candidate’s relevance, whereas the local SHAP scores measure the direction and magnitude of each biomarker candidate’s influence on the model output for one single pixel. SHAP maps deliver spatially localized explanations of the classification model’s decision-making process. Icons from [32, 33].

Traditionally, applications of supervised ML in IMS focus on maximizing the predictive performance of classification models designed to automate user-defined recognition tasks, without necessarily examining their decision-making processes. However, we suggest that examining the relationship between a classification model’s features and its prediction is important because it can reveal which features, and thus which molecular species, enable the differentiation of classes. Model interpretability is the ability to explain the predictions of a supervised ML model by reporting the relative predictive importance of its features 1. The importance, or relevance, of a feature is a measure of how it influences the model’s prediction, considering both its direct effect (i.e. statistical association with the prediction) and its indirect effect (i.e. statistical association between features) [24, 29]. The local predictive importance of a feature measures its influence on the predictive model’s output for a specific observation (e.g. the mass spectrum of one pixel), whereas the global predictive importance of a feature measures its influence on the predictive model’s output for all observations (e.g. all pixels of a sample) [24, 25, 29]. In addition to reporting which features drive the decision-making processes of supervised ML models, interpretability methods also facilitate model troubleshooting (e.g. debugging, monitoring, checking for bias). For example, in the context of IMS data analysis, interpretability methods make it possible to trace whether the decision-making process of a classification model is based on genuine biological patterns rather than on instrumental patterns or chemical noise that are spuriously associated to the class labels. ML interpretability methods effectively address the issue of supervised ML algorithms producing “black-box” models with unintelligible predictive mechanisms [23, 24, 25]. The importance of ML interpretability for knowledge discovery has recently been discussed in genomics [30] and single-cell mass spectrometry [31]. To our knowledge, our work is the first application of ML interpretability methods to IMS data for the purpose of biomarker candidate discovery. Our aim is to formulate and demonstrate how ML interpretability methods can be used to understand how the spatial distribution and relative abundance of certain molecular species relate to the classification of different regions of a tissue sample, effectively automating biomarker candidate discovery in IMS data.

Our approach to aiding biomarker discovery is to automate and accelerate the identification of promising biomarker candidates among discriminative molecular features (i.e. *m/z* values) by empirically learning which molecular species’ overexpression or underexpression enable the recognition of a user-defined class [20]. We translate the problem of biomarker discovery into a feature ranking problem: ML interpretability methods computationally estimate the importance of each feature with regards to a specific classification task and produce a ranking of the features in descending order of predictive importance. Ranking the features in terms of predictive importance facilitates the identification of a shortlist of molecular species that are characteristic of a class of interest, and thus have a higher likelihood of being useful biomarkers. In addition to providing one with a global understanding of which molecular species hold potential for recognizing a user-provided class, our approach uses SHAP maps to give the user spatially localized insight into each biomarker candidate’s relationship with the class of interest. SHAP maps are a novel graphical representation of a model’s decision-making process that can yield a nuanced local assessment of a biomarker candidate’s potential and spatial specificity. Our biomarker candidate discovery workflow is therefore a scalable computational tool that enables one to rapidly, efficiently, and automatically filter the multitude of molecular species recorded by IMS down to a panel of promising biomarker candidates that deserve further study and validation.

## 2 Machine learning methodology

### 2.1 Extreme gradient boosting for imaging mass spectrometry data classification

There are many applications of supervised ML in IMS: random forests [22, 34, 35], support vector machines [34, 36], convolutional neural networks [37, 38], and gradient boosting machines [39, 40] are frequently used classification model types. Decision trees are particularly suitable for IMS data analysis because they are non-linear and non-parametric predictive models that can account for complex dependencies between features, do not make assumptions about the underlying data distribution, and do not require feature scaling. A decision tree is a directed graph that partitions the feature space by recursive binary splitting: its nodes correspond to subsets of the data, and its branches correspond to the partitioning of a feature above or below a splitting threshold [41, 42, 43]. Given that a single decision tree is neither flexible nor stable enough to achieve high predictive performance on IMS data classification tasks, combining multiple decision trees into an ensemble model is usually a preferable strategy [42, 44]. We therefore choose to use extreme gradient boosting (XGBoost) models for classification. XGBoost is a fast and scalable implementation of (stochastic) regularized gradient boosting that was developed by Chen and Guestrin in 2016 [45] based on the work of Friedman [46, 47], Freund and Schapire [48]. An XGBoost model is an ensemble of regression trees (i.e. decision trees that output real values in their terminal nodes) that can perform classification by additive logistic modeling [49, 50].

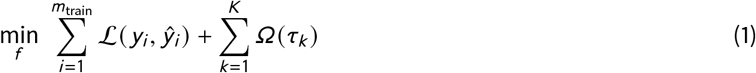

Regularized gradient boosting is a forward stagewise additive modeling algorithm for solving numerical optimization problems of the form of Equation 1. 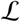 is a differentiable loss function (e.g. negative log-likelihood) that measures the difference between the observations’ labels *y_i_* and the predictive model’s predictions 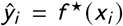, and *Ω* is a regularization term that penalizes the complexity of the regression trees making up the ensemble in order to avoid overfitting 2 [51, 45]. In Equation 1, the regression trees are written *τ_k_*, for *k* = 1, 2, 3…*K*, and *m*_train_ refers to the number of observations making up the training dataset. The XGBoost algorithm builds a classification model from sequentially added regression trees, each of which is focused on the observations that the previously added trees classified incorrectly [51, 52, 53]. Given an initial prediction *τ*_0_ (e.g. the logarithm of the odds), the accuracy of the ensemble model is iteratively improved by functional gradient descent: each newly added regression tree is parameterized to approximate the negative gradient of the loss function 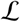 [51]. In order to avoid overfitting, the contribution of each newly added regression tree is weighted using a shrinkage parameter *ν*, with 0 < *ν* < 1 (*ν* = 0.3 in our case studies), which determines the learning rate of the boosting procedure [45, 54, 46]. In our automated biomarker candidate discovery workflow, the XGBoost learning process is stochastic because the regression trees making up the ensemble are learned on randomized subsamples of the training set, and because the features used for node splitting are chosen among a random subset of features [45, 54]. The idea is to randomly subsample the rows and columns of the data matrix during training in order to make each regression tree slightly different from the other regression trees, and hence prevent overfitting.

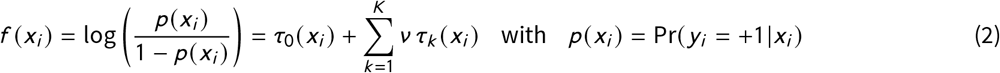

Equation 2 defines the raw output *f* (*x_i_*), or raw untransformed margin value, of an XGBoost classification model as the (natural) logarithm of the odds, called the log-odds [55, 54]. The odds are defined as the ratio of the probability *p* (*x_i_*) of observation *x_i_* being assigned to the positive class over the probability of observation *x_i_* being assigned to the negative class. The XGBoost classification model is an additive logistic regression model because it represents the log-odds as a linear combination of regression trees [52, 50]. The probability of the model predicting a positive outcome (i.e. assigning an observation to the class of interest) can be obtained from the log-odds thanks to a logistic transformation [50]: *p* (*x_i_*) = *S* (*f* (*x_i_*)) where *S* : ℝ ↦ [0, 1] is the sigmoid function. Since the sigmoid function is a non-decreasing saturation function, an increase in the log-odds implies an increase in the probability of predicting a positive outcome, and, conversely, a decrease in the log-odds implies a decrease in the probability of predicting a positive outcome. The classification model’s prediction 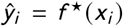 is either +1 or −1 depending on whether *p* (*x_i_*) is above or below a given threshold *η*, with 0 < *η* < 1 (*η* = 0.5 in our case studies).

### 2.2 Shapley additive explanations for measuring biomarker candidate relevance

Our workflow for biomarker candidate discovery in IMS data uses Shapley additive explanations (SHAP) to quantify the local and global predictive importance of features (e.g. *m/z* values in IMS) with respect to a given classification task. SHAP is a state-of-the-art interpretability method based on Shapley values from cooperative game theory. It regards the features as players that form coalitions (i.e. ordered subsets) to achieve the classification model’s output, which is the game’s payout. SHAP is called a model-agnostic interpretability method because it can derive post-hoc explanations for the predictions of any type of classification model by relating its input to its outputs [24, 25, 29]. SHAP was developed by Lundberg and Lee [56, 57] based on the work of Strumbelj and Kononenko [58, 59], and on Ribeiro *et al.*’s idea of locally approximating the decision-making process of a “black-box” supervised ML model using inherently interpretable local surrogate models [60].

In order to explain the prediction made by a classification model on a specific observation, SHAP computes the contribution of each feature to the model’s output using Shapley values. The Shapley value of a feature is its contribution to the model’s output for a specific observation, averaged over all possible feature orderings [57, 61]. In the words of Lundberg *et al.*, “Shapley values are computed by introducing each feature, one at a time, into a conditional expectation function of the model’s output, and attributing the change produced at each step to the feature that was introduced” [61]. Equation 3 defines the Shapley value 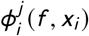 of feature *x ^j^*, with *j* ∈ {1, 2, 3…*n*}, when explaining the predictive model’s decision-making process for one specific observation *x_i_* = *X*_(*i*,:)_, with *i* ∈ {1, 2, 3…*m*}. Since a feature’s contribution to the model’s output depends on the order in which other features were introduced, the feature’s Shapley value is obtained by averaging its contribution over all possible feature orderings. In Equation 3, the set of all possible feature orderings is written *Π*. The set of features that we are conditioning on, written 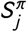, is the set of all features that precede feature *x ^j^* in ordering *π*.

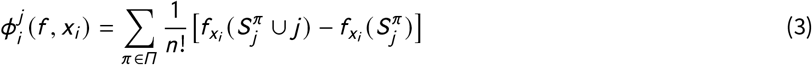

The set function 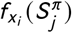 is defined by Equation 4 as the conditional expectation of the classification model’s output. The *n*-dimensional vector *x_i_* is considered to be a random variable where only the features belonging to subset 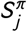 (i.e. the features before *j* in the feature ordering *π*) are known [62]. The unknown features (i.e. the features after *j* in the feature ordering *π*) are obtained by sampling from the training dataset [62]. Note that *f* (*x_i_*) in Equation 4 is the model’s raw output for observation *x_i_*, rather than the predicted class label *f* ^⋆^(*x_i_*) ∈ {−1, +1}. By focusing on the raw model output, SHAP does not require knowledge of an observation’s true class label (*y_i_* for *i* ∈ 1, 2, 3…*m*) to evaluate the degree to which a classification model depends on a specific feature. SHAP can therefore be used to explain the decision-making process of a model on new unlabeled data, which is useful for measuring the influence of each feature on the model’s generalization performance.

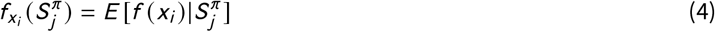

The sign of a feature’s Shapley value provides information about the direction of its effect on a classification model’s output. A positive Shapley value indicates that feature *x ^j^* increases the raw output *f* (*x_i_*) of the predictive model for observation *x_i_*. Conversely, a negative Shapley value indicates that feature *x ^j^* decreases the raw output. The Shapley value’s magnitude indicates how strongly the corresponding feature influences the classification model’s local decision-making process. In our work, we refer to the Shapley value of a feature for a given observation as its local SHAP importance score. In the context of IMS data classification, the Shapley value 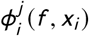 reports the contribution of the *j* ^th^ m/z bin or peak when assigning the *i* ^th^ pixel’s mass spectrum to a class. Computing the local SHAP importance scores of all features (i.e. *m/z* values) for all observations (i.e. mass spectra) yields an *m* × *n* matrix whose (*i*, *j*)^th^ entry is the Shapley value of feature *x ^j^* for observation *x_i_*.

SHAP owes its reliability to the fact it satisfies the local accuracy and consistency properties [61]. The local accuracy property, also known as the efficiency property in cooperative game theory, guarantees that the Shapley values of all features add up to the difference between the predictive model’s raw output *f* (*x_i_*) for a given observation *x_i_* and the model’s expected output *E* [*f* (*x_i_*)] over the entire dataset [61]. The local accuracy property is given by Equation 5. SHAP offers contrastive explanations that compare the model’s local output to it’s average global output. In IMS terminology, the local accuracy property states that, given a mass spectrum of interest, the sum of the Shapley values of its molecular features (i.e. *m/z* values) is equal to the classification model’s raw output for that mass spectrum minus the model’s average raw output over all mass spectra. SHAP distributes the difference between the model’s output for a mass spectrum of interest and the model’s average output, among the different *m/z* values that the model uses as inputs.

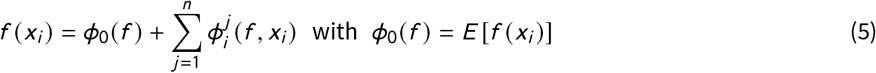

The consistency property, also known as the monotonicity property in cooperative game theory, states that if a classification model changes so that some feature’s influence on the output increases, the importance score assigned to that feature does not decrease [63]. Consistency is necessary for the ranking of a model’s features according to their importance scores because it guarantees that a feature with a higher importance score than another feature is actually more important to the model than the other feature. Note that impurity-based measures of global feature importance, which are popular for measuring feature importance in decision tree ensembles 3 and have been used in IMS [39], are actually inconsistent and can therefore produce unreliable feature rankings [63].

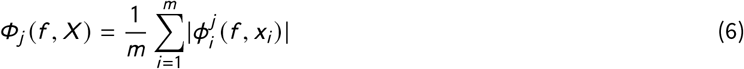

A global measure of feature importance can be obtained by averaging the magnitude of each feature’s local SHAP scores, or Shapley values, over all observations in the dataset [61]. Equation 6 defines what we refer to as the global SHAP score *ϕ_j_* of feature *x ^j^* = *X*_(:,*j*)_ for *j* ∈ {1, 2, 3…*n*}. The global SHAP score of a feature quantifies its influence on the model’s decision-making process, averaged over all possible feature orderings and all observations. Computing the global SHAP importance scores of all features yields an *n*-dimensional vector where *n* is the total number of features. In the context of IMS data analysis, the global SHAP score of a feature is an experiment-wide measure of the feature’s predictive importance with respect to a given classification task. Promising biomarker candidates can be easily identified by ranking IMS features (i.e. *m/z* values) in descending order of global SHAP importance. Retaining the top-ranking features yields a shortlist of biomarker candidates that are worthy of further study.

In our workflow for biomarker candidate discovery in IMS data, we use a fast implementation of SHAP called TreeSHAP, or TreeExplainer [67], that is specific to decision tree based predictive models like XGBoost. Unlike other SHAP implementations (e.g. KernelSHAP) that calculate sampling-based approximations of Shapley values (often in exponential time), TreeSHAP is able to compute the exact Shapley values of features within low-order polynomial time by exploiting the structure of decision trees [63, 61]. When using TreeSHAP to measure the local and global SHAP importance scores of features, one has to choose between two feature perturbation approaches [67]. In this paper, we opt for the tree-path dependent approach because it involves computing the observational, rather than interventional, Shapley values [62]. Observational Shapley values are defined by Equations 3 and 4, whereas interventional Shapley values define the set function differently. The difference between observational and interventional Shapley values relates to how SHAP handles statistical dependencies between the features that the model uses as inputs [62]. Accounting for high-dimensional feature dependencies is what makes measuring the predictive importance of IMS features, many of which are involved in common biochemical pathways, particularly challenging. Another measure of global feature predictive importance, called permutation importance (PI), has been used for ranking IMS features with regards to tissue classification tasks [22] despite it only partially accounting for feature inter-dependencies. PI is a popular global model-agnostic interpretability method that defines the importance of a feature as the average decrease in model accuracy when its values are randomly permuted across all observations4. PI only accounts for correlation, whereas SHAP also accounts for high-order feature inter-dependencies and enables the discovery of both linear and non-linear patterns in the data [61]. Furthermore, PI relies upon out-of-distribution data instances (that are not necessarily realistic), whereas computing the global SHAP score of a feature using observational Shapley values constrains the sampling of unknown features to a range of values (i.e. partitions of the feature space) allowed by the decision trees making up the ensemble [69]. A detailed discussion of how TreeSHAP computes observational Shapley values, and how observational Shapley values handle feature dependencies, is beyond the scope of this paper, and we therefore refer the reader to [61, 62, 63]. Observational Shapley values are recommended for knowledge discovery in biology and chemistry because they spread credit among correlated features that are jointly informative of the outcome of interest [62].

### 2.3 SHAP maps for a spatial understanding of a classification model’s decision-making process

In addition to automatically establishing an experiment-wide biomarker candidate shortlist by means of global SHAP score ranking, we furthermore introduce a novel spatially-aware representation of local SHAP-based explanations, called a SHAP map. The SHAP map of a molecular feature is obtained by plotting that feature’s local SHAP importance scores, or Shapley values, across all pixels. SHAP maps facilitate a spatially localized understanding of a classification model’s decision-making process. In the context of biomarker candidate discovery, SHAP maps provide one with a nuanced and location-specific (e.g. cell type specific, tissue region specific) view into a molecular species’ biomarker potential. Unlike global SHAP importance scores, local SHAP importance scores avoid conflating the magnitude of the feature’s effect with the prevalence of its effect across the sample surface area.

The SHAP map of a feature answers the following two questions:

**• Where does the feature increase or decrease the classification model’s output?**

The feature increases the probability of the model assigning a pixel to the class of interest (i.e. the positive class) where its local SHAP scores are positive (red pixels). The feature decreases the probability of the classification model assigning a pixel to the positive class where its local SHAP scores are negative (blue pixels). In our application, studying the sign of a feature’s Shapley values together with the feature’s spatial distribution (e.g. the feature’s ion image) enables the user to determine whether it is the presence or the absence of a feature that is indicative of the biological state or disease condition of interest. If the regions where the feature’s measured intensity is high coincide with the regions where the feature’s Shapley values are positive, the feature’s presence is indicative of the class of interest. The relationship between the feature’s abundance and the class prediction is correlative. Conversely, if the regions where the feature’s measured intensity is low coincide with the regions where the feature’s Shapley values are positive, the feature’s absence is indicative of the class of interest. The relationship between the feature’s abundance and the class prediction is anticorrelative.

**• Where does the feature strongly or weakly influence the classification model’s output?**

The feature has a relatively large influence on the classification model where its Shapley values have a high magnitude (pixels with high saturation). Conversely, the feature has a relatively small influence on the model where its Shapley values have a low magnitude (pixels with low saturation). Studying the magnitude of a feature’s Shapley values provides insight into how large or small the feature’s local influence on a model is. In our application, we consider a feature (i.e. *m/z* value) to be relevant to recognizing a given class in the regions of the sample were its Shapley values have a high magnitude.

## 3 Results & Discussion

Our biomarker candidate discovery workflow is demonstrated on two IMS datasets that were acquired by MALDI quadrupole time-of-flight (Q-TOF) IMS using the prototype MALDI timsToF Pro (Bruker Daltonics, Germany) in positive ion mode [70]. Please refer to the supplementary material for information regarding the materials, sample preparation, experiments, histology, and IMS data preprocessing. Since the following five case studies do not involve the study of diseased tissue, the ranked features are not indicative of any pathological processes but rather of anatomical structures. Therefore, the term “molecular marker” is preferred over the term “biomarker” in Section 3. It should be noted that, methodologically speaking, there is no difference: in both cases our workflow looks for differentiating markers (i.e. *m/z* values) corresponding to user-provided classes of interest.

- Dataset n°1 was acquired from the sagittal whole-body section of a mouse-pup. The autofluorescence microscopy image of the tissue section is presented in Figure 3a and was used to guide annotation of the regions of interest [71]. The sample was cryosectioned at 20 μm thickness and a 1,5-diaminonaphthalene matrix was applied by sublimation. The mean mass spectrum of the dataset was retrieved and peak-picked to produce a feature list of 879 distinct ion species. The *m/z* acquisition range is 300-1,200 and the pixel size is 50 μm×50 μm. The dataset consists of a total of 164,808 pixels. Our workflow is therefore applied to a dataset of 164,808 observations and 879 features. The challenge of molecular marker discovery in the two case studies tied to this dataset therefore amounts to automatically determining which molecular species, among the 879 measured *m/z* values, are most relevant to recognizing two anatomical regions: the mouse-pup’s brain and its liver.
- Dataset n°2 was acquired from the sagittal section of a rat kidney. The hematoxylin & eosin stained microscopy image of the tissue section is presented in Figure 3b. The sample was cryosectioned at 12 μm thickness and a 1,5-diaminonaphthalene matrix was applied by sublimation. The mean mass spectrum of the dataset was retrieved and peak-picked to produce a feature list of 1,428 distinct ion species. The *m/z* acquisition range is 300-2,000 and the pixel size is 15 μm×15 μm. Our workflow is applied to a data table of 591,534 observations and 1,428 features. The challenge of molecular marker discovery amounts to automatically determining which molecular species, among the 1,428 measured m/z values, are most relevant for recognizing three different regions of the kidney: the cortex, the inner and outer medulla.

**Figure 3.**
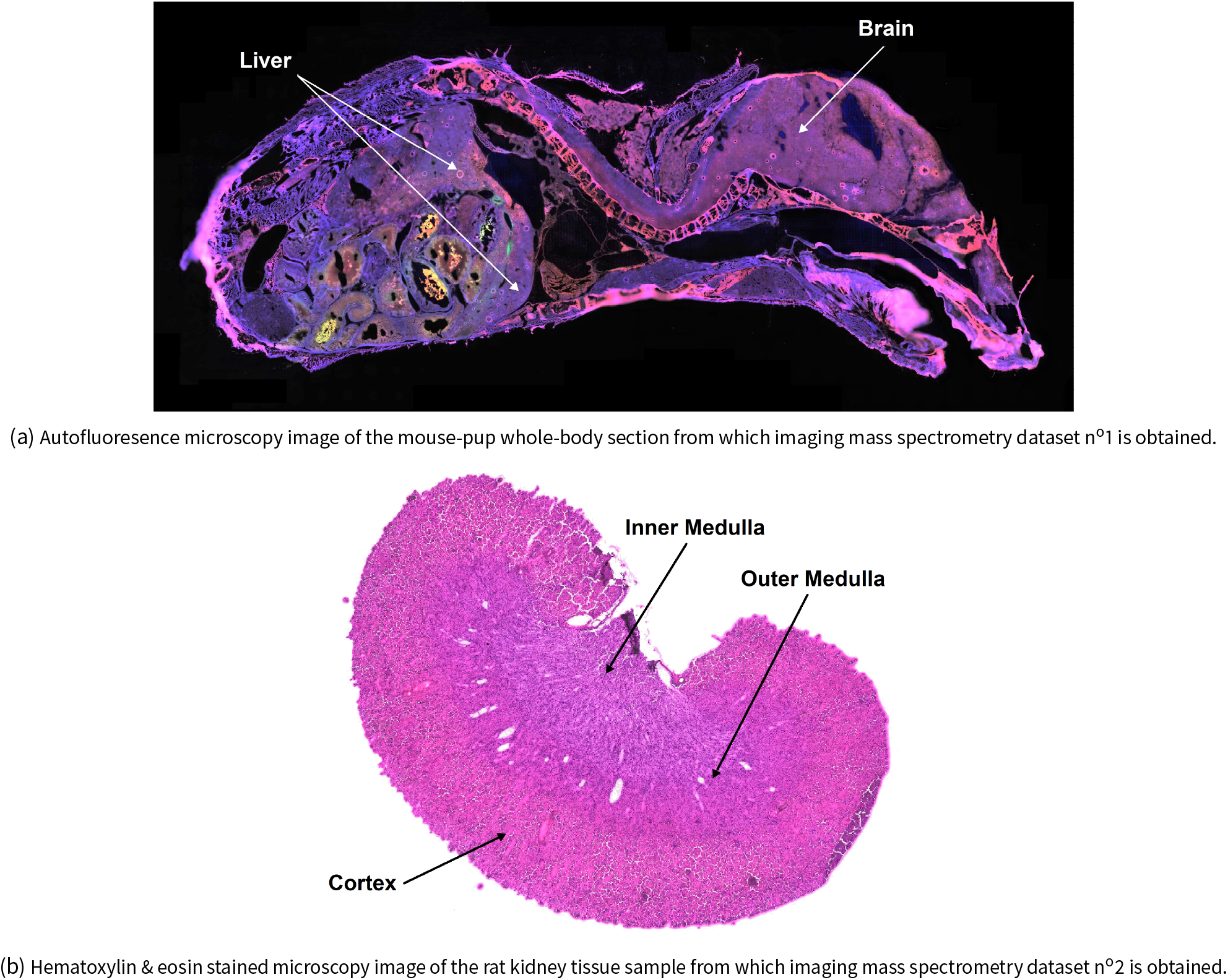
Microscopy images of the tissue sections imaged in IMS datasets n°1 and n°2.

Five anatomical regions were delineated within the two tissue samples on the basis of the microscopy images in Figure 3. Each tissue region was given a class label: a brain and a liver region in dataset n°1 and a cortex, inner medulla, and outer medulla region in dataset n°2. Our aim is to discover molecular markers for each of these user-provided class labels. The molecular marker discovery is treated separately for each class, using the one-versus-all procedure, yielding five binary classification problems whose target (i.e. positively labeled) classes are the following: the mouse-pup’s brain and liver in dataset n°1; the rat kidney’s inner medulla, outer medulla, and cortex in dataset n°2. Note that, although user-defined masks are employed to label the data in our case studies, our approach would work equally well if provided with automatically-generated class annotations. The first three case studies (i.e. discovering molecular markers for the mouse-pup’s brain and liver in dataset n°1, discovering molecular markers for the rat kidney’s inner medulla in dataset n°2) are covered in Section 3, whereas the remaining two case studies (i.e. discovering molecular markers for the rat kidney’s outer medulla and cortex in dataset n°2) are provided in the supplementary material.

As discussed in subsection 2.1, XGBoost models are used to classify the pixels on the basis of their mass spectra. These five classification problems are imbalanced because their corresponding datasets have unequal class cardinality (i.e. the negatively labeled pixels outnumber the positively labeled pixels). We avoid using accuracy (i.e. the proportion of predictions that are correct) to measure the classification models’ predictive performance since accuracy tells us little about whether false negatives or false positives are more common [72]. Instead, we choose to measure our classification models’ predictive performance using balanced accuracy, precision, and recall. Recall (also called sensitivity or the true positive rate) is the proportion of positive observations that are correctly identified. Precision is the proportion of all positive predictions that are correct. Specificity (also called the true negative rate) is the proportion of negative observations that are correctly identified. Balanced accuracy is the arithmetic mean of sensitivity and specificity [72].

As discussed in subsections 2.2 and 2.3, the TreeSHAP implementation of the SHAP interpretability method (with observational Shapley values) is used to rank the features (i.e. *m/z* values) in descending order of global predictive importance. The top-ranking features are highly discriminative with regards to a labeled tissue class and are therefore considered to be promising molecular markers for that class of interest. In addition to automatically establishing a shortlist of molecular species that are statistically related to user-provided tissue class labels, our workflow delivers spatially localized insight into the relationship (e.g. correlative, anticorrelative) between each measured ion species and the class of interest by means of a novel visualization approach called SHAP maps.

### 3.1 Dataset n°1: Recognition of the brain and liver of a mouse-pup

Classification-oriented supervised ML algorithms require labeled training data (in our case, labeled pixels) to build a classification model. In the two mouse-pup case studies, anatomical class labels are obtained as user-provided spatial delineations of the mouse-pup’s brain and liver in the tissue sample. Exploratory analysis of the IMS data was performed using non-negative matrix factorization to aid in that delineation task [73, 74, 21]. The low-dimensional latent patterns extracted by non-negative matrix factorization from dataset n°1 directly delineated several organs within the mouse-pup whole-body section, which facilitated easier and more robust manual localization and annotation of the target organs. In the context of molecular marker discovery, the target organs (or tissue regions, cell types, or cells) that are provided as masks to the supervised ML algorithm are also the organs (or tissue regions, cell types, or cells) for which we want to discover molecular markers. Figure 4 shows a spatial representation of the masks used to build the XGBoost classification models for the mouse-pup cases. Pixels are either annotated as belonging to the target organ (i.e. positive class) or not belonging to the target organ (i.e. negative class). Some pixels (e.g. at the borders of target organs) were difficult to annotate definitively and were excluded from the training set to avoid providing the supervised ML algorithm erroneous or unreliable training examples. Furthermore, the negative class was downsampled to avoid the one-versus-all classification of the brain and liver being severely imbalanced. After down-sampling, ~25% of the pixels used to build the model belong to the positive class, and ~75% belong to the negative class.

**Figure 4.**
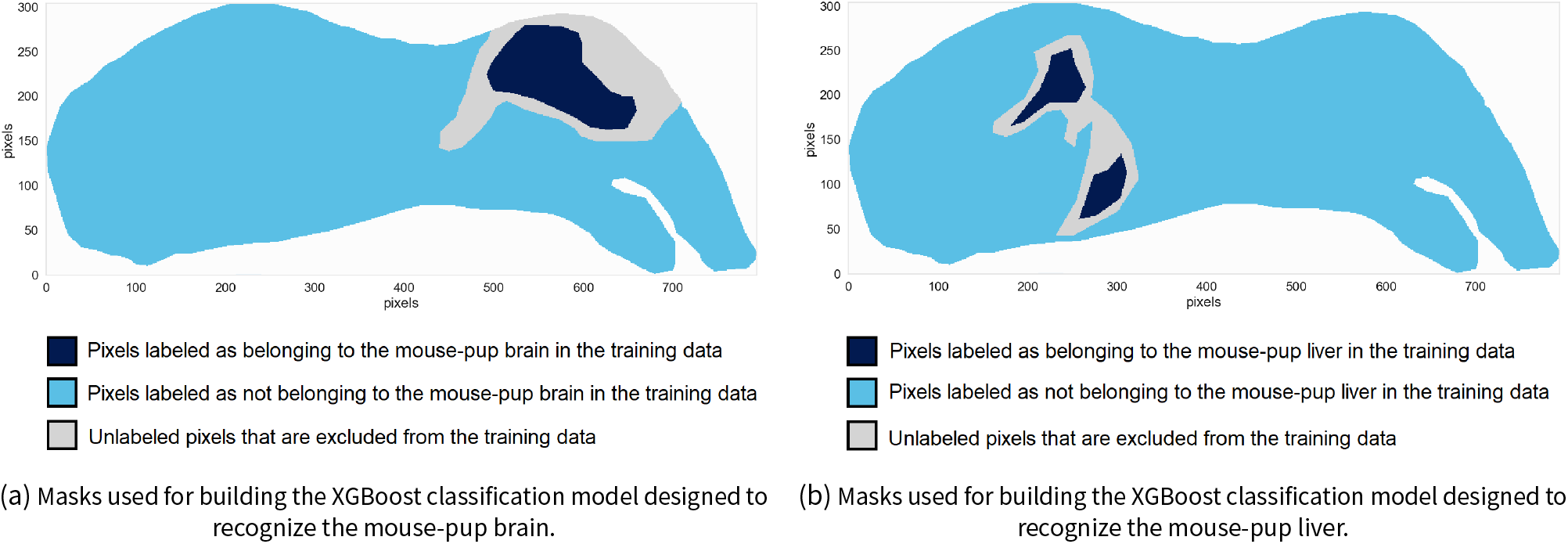
Class-defining masks used as inputs for training the two XGBoost classification models designed to recognize the mouse-pup brain and liver. For each task, regions of the tissue sample were manually annotated as belonging to one of three categories: dark blue pixels are labeled as belonging to the target organ and make up the positive class, light blue pixels are labeled as not belonging to the target organ and make up the negative class, and gray pixels are close to borders between the target organ and other anatomical structures, making it difficult to annotate them definitively. The latter are therefore excluded from the training data to avoid feeding the model unreliable annotations during training.

#### Molecular marker discovery for the mouse-pup brain

Our brain molecular marker discovery workflow starts with building a classification model from IMS dataset n°1 and the user-provided brain mask shown in Figure 4a. The model building process is illustrated in Figure 1a. We obtain an XGBoost model that automatically recognizes brain tissue pixels on the basis of their mass spectra. To demonstrate the recognition capabilities of the learned XGBoost model, we supply it with all IMS measurements (both labeled and unlabeled pixels), effectively going through the prediction process illustrated in Figure 1b. Figure 5a, which is the result of the prediction process, shows which mouse-pup tissue regions are predicted to belong to the brain according to the XGBoost classification model. As is apparent from Figure 5a, the mouse-pup’s brain (as well as parts of its spinal cord) are successfully differentiated from the other organs. Regarding generalization performance, the XGBoost model trained to recognize pixels belonging to the mouse-pup brain achieves a balanced accuracy of 0.9925, a precision of 0.9967, and a recall of 0.9974. Note that generalization performance is evaluated on a testing dataset, which is a labeled selection of pixel measurements that differ from the training dataset.

**Figure 5.**
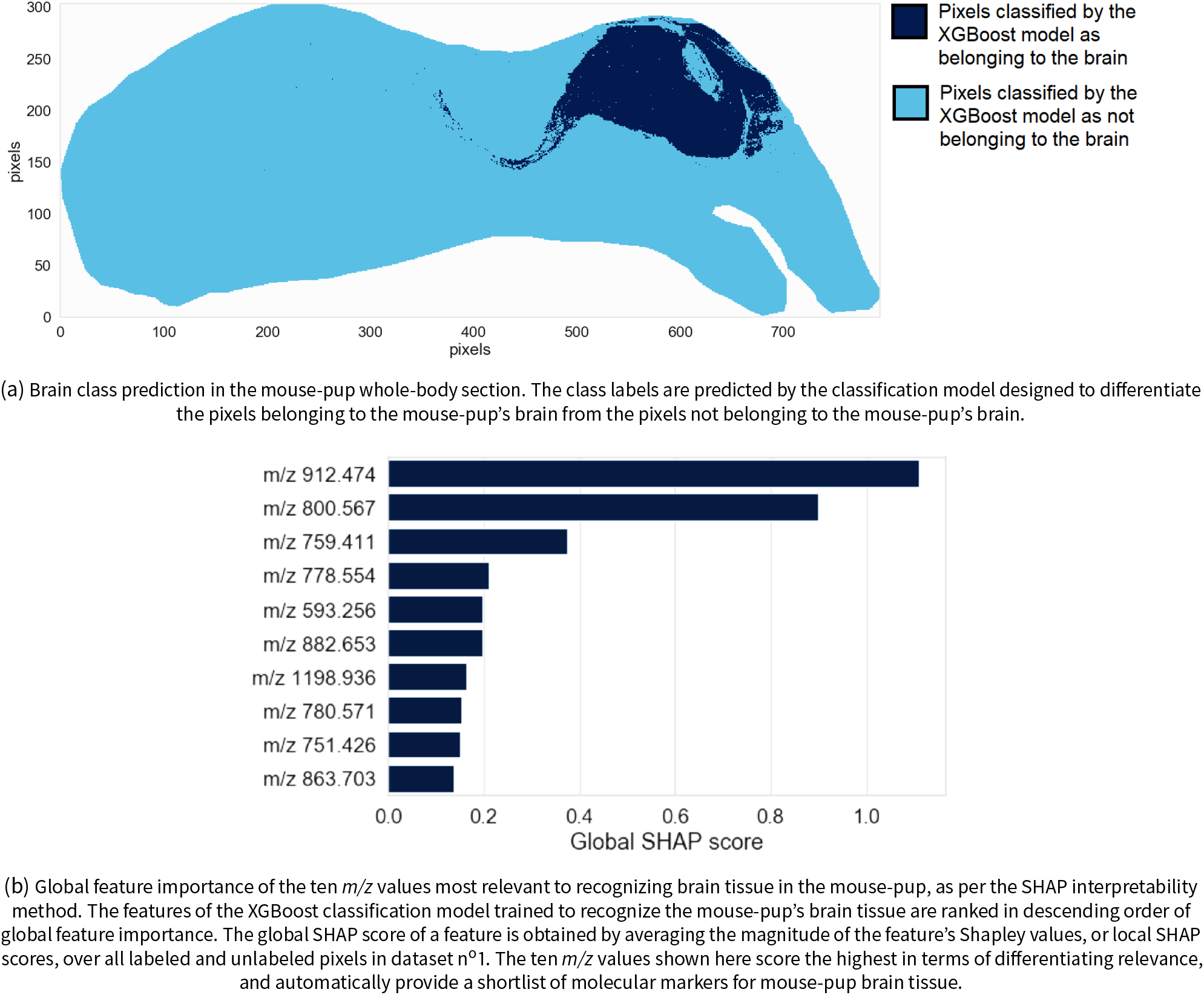
Mouse-pup brain recognition and global feature ranking.

Figure 5b shows the top ten molecular markers of the global ranking of 879 features (i.e. *m/z* values) obtained by TreeSHAP. The features are ranked in descending order of global SHAP score, and thus in descending order of relevance to brain tissue recognition, yielding a shortlist of molecular markers for mouse-pup brain tissue. Figure 5b provides insight into a feature’s global (i.e. tissue-wide) relevance to the recognition task of Figure 5a. However, the spatially localized nature of IMS measurements together with the SHAP map representation developed above allows us to obtain tissue location specific insights into an *m/z* value’s relevance. Figure 6 shows the ion images and SHAP maps of the three top-ranking features of Figure 5b. The left column of Figure 6 displays the spatial distribution and relative abundance of the three top-ranking molecular features for recognizing the mouse brain. Figures 6a, 6c, and 6e are ion images of the features ranked n°1, n°2, and n°3 respectively, and they are displayed using a pseudo-color scale whose brightness is indicative of the signal intensity measured at a given pixel. These ion images provide a classical view on molecular distribution by reporting the ion intensity signal corresponding to the molecular species at hand.

**Figure 6.**
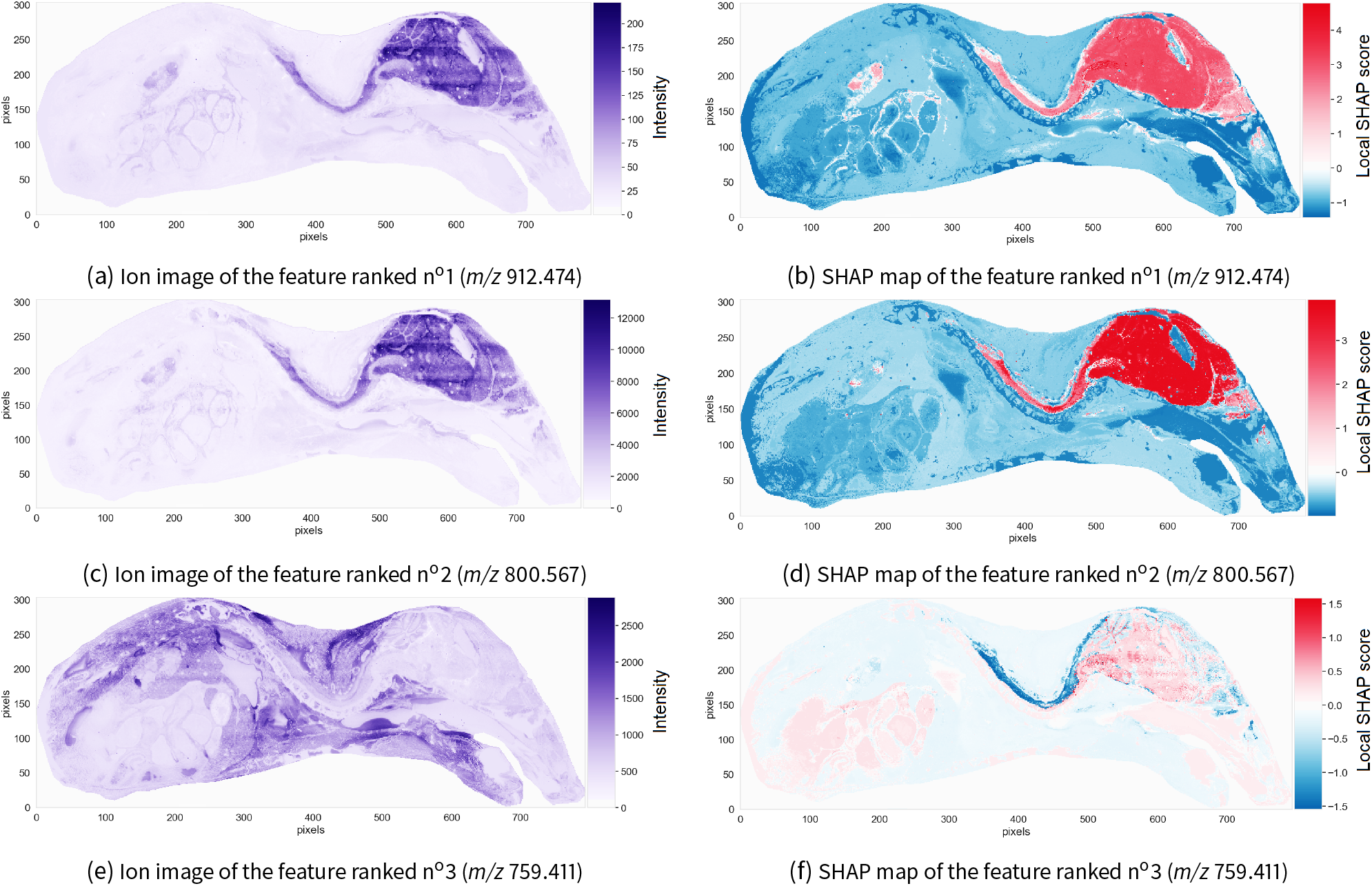
Three promising molecular markers for the mouse-pup’s brain. The ion images (left) and SHAP maps (right) of three features (i.e. *m/z* values) with the most influence on the decision-making process of the classification model trained to recognize the mouse-pup’s brain are shown. The ion images plot the spatial distribution and measured intensity of each feature across the sample, and are not specifically tied to the task of recognizing the brain. The SHAP maps plot the spatial distribution of Shapley values, or local SHAP predictive importance scores, of each feature across the sample, and provide information on where and how the feature is relevant to the task of recognizing brain.

However, ion images do not provide any information about how that ion intensity relates to the recognition of brain tissue. The right column of Figure 6 provides information on the signs and magnitudes of the local SHAP scores across the sample for each top-ranking feature. Figures 6b, 6d, and 6f are the SHAP maps of the features ranked n°1, n°2, and n°3 respectively. These SHAP maps provide information on where and how a given ion intensity signal relates to the task of brain tissue recognition.

Figure 6a is the ion image of the feature ranked n°1, whose *m/z* value is 912.474. The measured intensity of this feature is high in the brain and spinal cord. According to Figure 6b, feature n°1 increases the log-odds (raw) output of the XGBoost model in the brain region: the Shapley values in the brain and spinal cord are positive, and negative elsewhere. The presence of feature n°1 increases the log-odds (and probability) of the XGBoost classification model predicting that a given pixel belongs to the brain. The ion image and SHAP map of the feature ranked n°2 (*m/z* 800.567) are very similar to those of the feature in first position. Both top-ranking features are positively correlated with the model assigning a pixel to the brain. Measuring high intensity signals for features n°1 and n°2 in a given pixel increases the log-odds (and probability) of the model assigning that pixel to the brain. Given the high predictive performance of the XGBoost classification model, which indicates that the model is probably a good approximation of the biochemical processes taking place in the tissue, it can be assumed and inferred that measuring a high intensity signal for features n°1 and n°2 in a given pixel also increases the probability of that pixel actually belonging to the brain. In other words, the presence of these features (*m/z* values) is characteristic of the mouse-pup’s brain and spinal cord and differentiates the brain and spinal cord from other regions in the tissue.

Figure 6e indicates that the feature ranked n°3, whose *m/z* value is 759.411, has a low intensity both in the brain and the spinal cord. Its measured intensity in the spinal cord is slightly higher than its intensity in the brain. Figure 6f shows that the Shapley values of that feature are negative in the spinal cord (with a magnitude between −1.0 and −1.5). The area highlighted (negatively, hence in dark blue) in Figure 6f, namely the spinal cord, is where feature n°3 plays a role in helping to obtain a biomolecular signature unique to the brain. The way in which this feature helps the classification model correctly identify the brain pixels can be read from the sign of its Shapley values, or local SHAP scores. The local SHAP values in the spinal cord are negative, meaning that whatever the signal is that is measured for this feature in the spinal cord, it lowers the log-odds (and probability) of assigning a pixel to the brain. Studying the ion image of feature n°3 furthermore reveals that the ion intensity for *m/z* 759.411 is low in the spinal cord, but still higher than in the brain. This means that a relative increase in signal intensity of *m/z* 759.411 strongly decreases the log-odds (and probability) of predicting a pixel belonging to the brain. Unlike the features ranked n°1 and n°2 that are good molecular markers for both the brain and spinal cord, the feature ranked n°3 enables the XGBoost classification model to tell the brain apart from the spinal cord. We would not be able to differentiate the mouse’s brain from its spinal cord if we were to use only the two top-ranking features (*m/z* 912.474 and *m/z* 800.567). This example illustrates the subtle understanding of molecular marker spatial specificity that can be obtained from SHAP maps. If one needs a molecular marker for both the brain and spinal cord, both *m/z* 912.474 and *m/z* 800.567 are good candidates. If one requires the ability to tell brain tissue apart from spinal cord tissue, a more elaborate panel of molecular markers is proposed: if *m/z* 912.474 and *m/z* 800.567 are present in high abundance in a tissue area, and if *m/z* 759.411 is present in very low abundance, the probability of those pixels describing brain tissue (exclusively) is very high.

#### Molecular marker discovery for the mouse-pup liver

Our liver molecular marker discovery workflow starts with building a classification model from IMS dataset n°1 and the user-provided liver mask shown in Figure 4b. Figure 7a shows the result of applying the classification model to all pixels of dataset n°1: the mouse-pup’s liver is successfully differentiated from the other organs, demonstrating that the model was able to capture a liver-specific mass spectral signature in the data. Regarding generalization performance, the XGBoost classification model trained to recognize pixels belonging to the mouse-pup liver achieves a balanced accuracy of 0.9960, a precision of 0.9984, and a recall of 0.9989.

**Figure 7.**
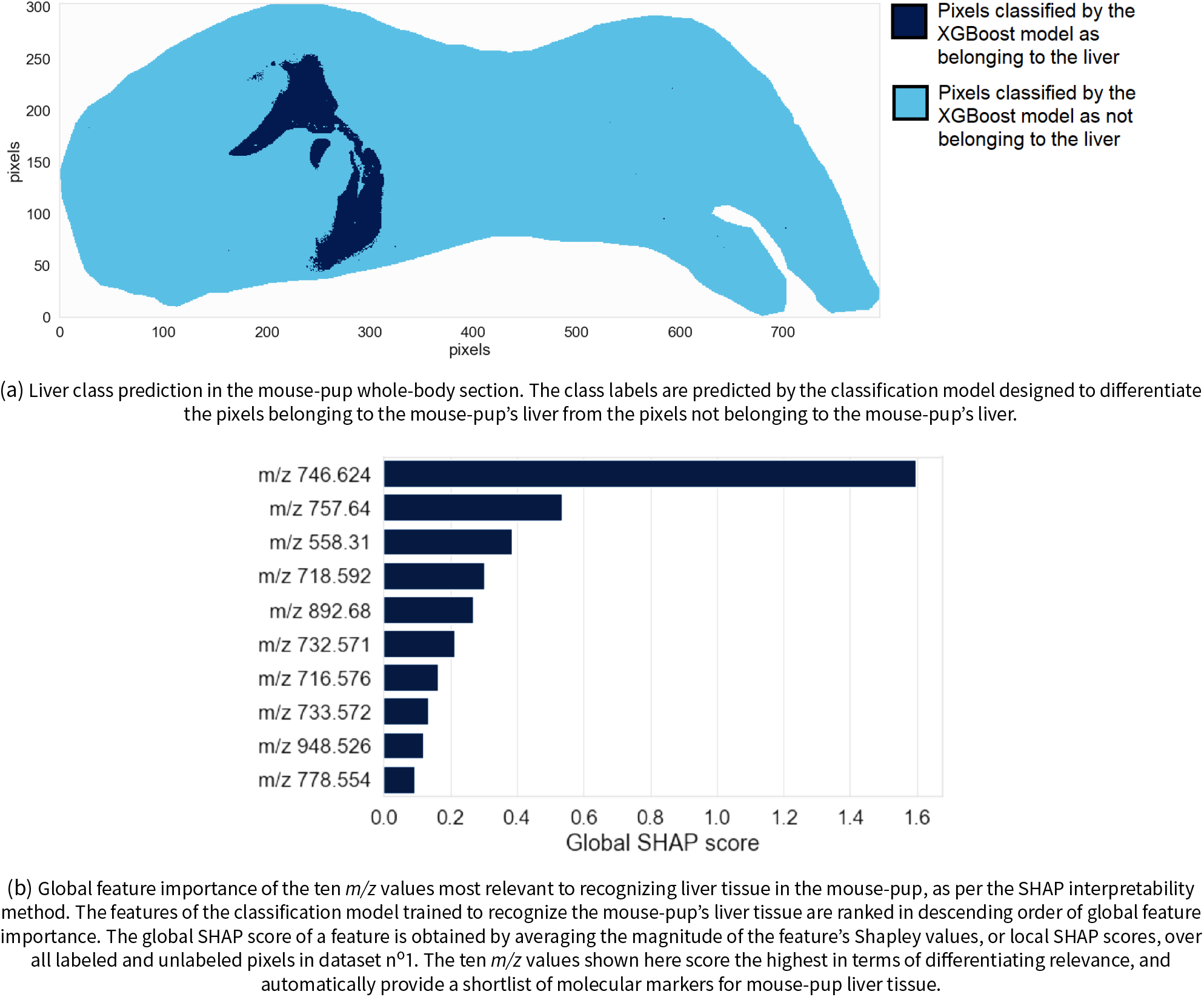
Mouse-pup liver recognition and global feature ranking.

Figure 7b shows the top ten molecular markers out of the global ranking of 879 features (i.e. *m/z* values) as obtained by TreeSHAP. The features are ranked in descending order of global SHAP predictive importance score, and thus in descending order of relevance to liver tissue recognition. Figure 7b therefore provides insight into a feature’s global relevance to the recognition task of Figure 7a. The feature with the highest influence on the XGBoost classification model used to assign pixels to the liver (or not) has a *m/z* value of 746.624 and a global SHAP score of 1.595.

The left column of Figure 8 provides information about the spatial distribution and relative abundance of the three top-ranking molecular features for recognizing the mouse liver in this dataset. Figures 8a, 8c, and 8e are the ion images of the features ranked n°1, °2, and °3 respectively. The right column of Figure 8 provides information on the signs and magnitudes of the local SHAP scores, or Shapley values, across the sample for each top-ranking feature. Figures 8b, 8d, and 8f are the SHAP maps of the three top-ranking features. These SHAP maps provide information on where and how a given ion intensity signal relates to the task of liver tissue recognition. Studying the left and the right columns of Figure 8 enables one to understand which features are positively or negatively correlated with a pixel being assigned to the liver.

**Figure 8.**
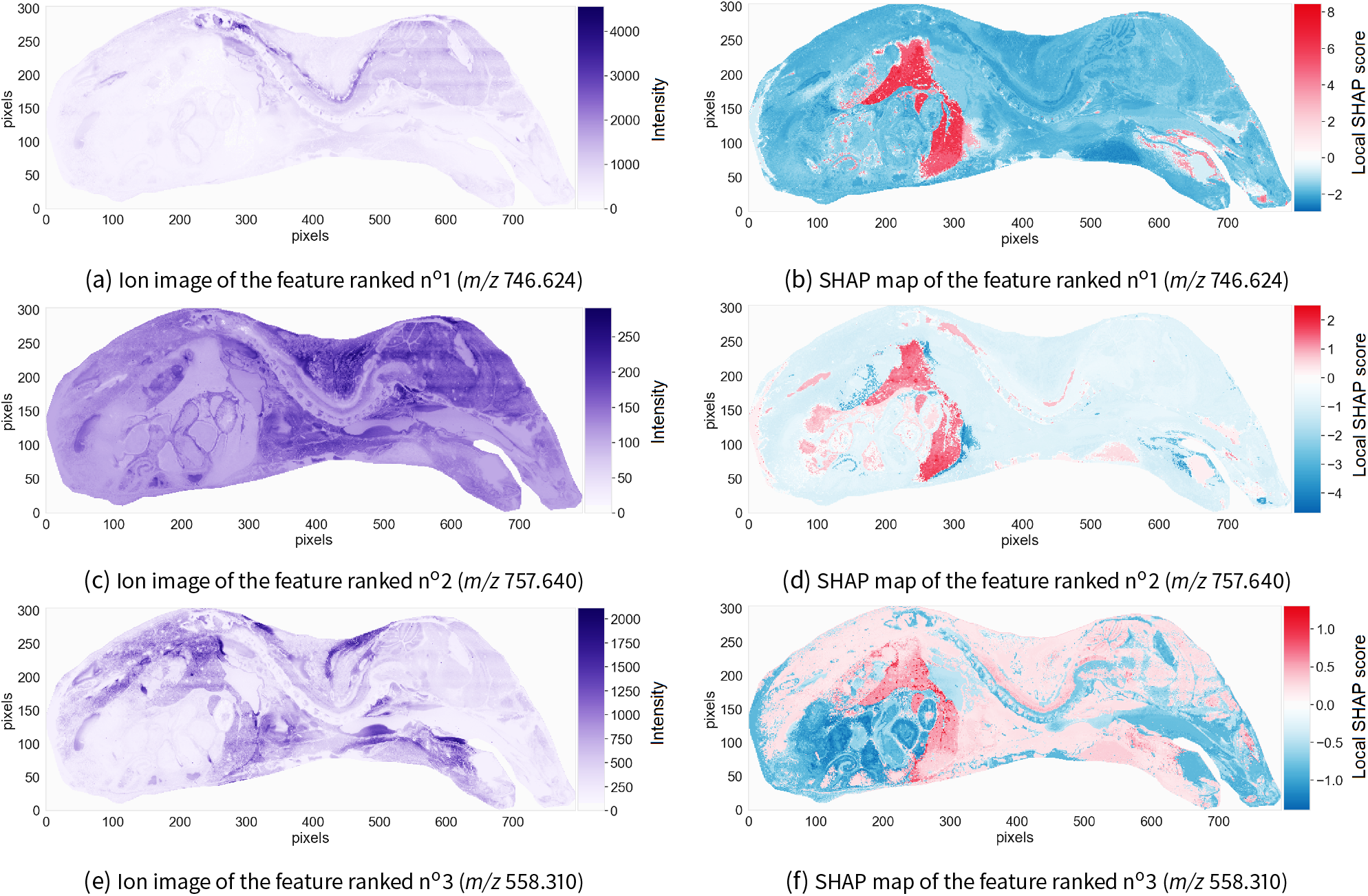
Three promising molecular markers for the mouse-pup’s liver. The ion images (left) and SHAP maps (right) of three features (i.e. *m/z* values) with the most influence on the decision-making process of the classification model trained to recognize the mouse-pup’s liver are shown. The ion images plot the spatial distribution and measured intensity of each feature across the sample, and are not specifically tied to the task of recognizing the liver. The SHAP maps plot the spatial distribution of Shapley values, or local SHAP predictive importance scores, of each feature across the sample, and provide information on where and how the feature is relevant to the task of recognizing the liver.

The ion image of the feature ranked n°1 (Figure 8a) indicates that it has a very low abundance in the liver region. Yet the SHAP map of that feature (Figure 8b) shows that the local SHAP scores are positive, with a high magnitude, in the liver region. The feature ranked n°1 can therefore be considered negatively correlated with a pixel being assigned to the liver. The feature ranked n°2 is also negatively correlated with the XGBoost classification model assigning a pixel to the liver. In short, measuring a low intensity for features ranked n°1 and n°2 in a given pixel increases the log-odds (and probability) of the classification model assigning that pixel to the liver. Given the high predictive performance of the classification model, we can assume that measuring a low intensity of features ranked n°1 and n°2 in a given pixel also increases the probability of that pixel actually belonging to the liver. Conversely, the feature ranked n°3 is positively correlated with a pixel being assigned to the liver: its ion image (Figure 8e) indicates that the feature ranked n°3 has a high abundance in the liver, and its SHAP map (Figure 8f) indicates that its local SHAP scores are high in the liver region. Measuring a high intensity of the feature ranked n°3 in a given pixel increases the log-odds (and probability) of the classification model assigning that pixel to the liver. Furthermore, given the high predictive performance of the classification model, we can assume that measuring a high intensity of the feature ranked n°3 in a given pixel also increases the probability of that pixel actually belonging to the liver. The absence of features ranked n°1 and n°2, and the presence of the feature ranked n°3 are therefore characteristic of the mouse-pup liver. If one measures a low abundance of *m/z* 746.624 and *m/z* 757.640 and a high abundance of *m/z* 558.310 in a pixel, the probability of that pixel belonging to the liver is very high.

### 3.2 Dataset n°2: Recognition of renal inner medulla, outer medulla, and cortex

Annotating the three different functional tissue regions of the rat kidney - namely the inner medulla, outer medulla and cortex - is required to generate the class labels needed to train the three corresponding XGBoost classification models. Similar to the previous case studies (subsection 3.1), exploratory analysis by means of non-negative matrix factorization was used to aid in delineating masks. Figure 9 shows the pixels annotated as belonging to one of the three target regions. The pixels that were difficult to annotate manually were excluded from the training and testing datasets. Similar to the previous case studies (subsection 3.1), downsampling of the negative class was performed. The inner medulla, outer medulla, and cortex are differentiated from the other two regions using one-versus-all classification where the target region is the positive class, and the two other regions make up the negative class. Please refer to the supplementary material for the outer medulla and cortex case studies.

**Figure 9.**
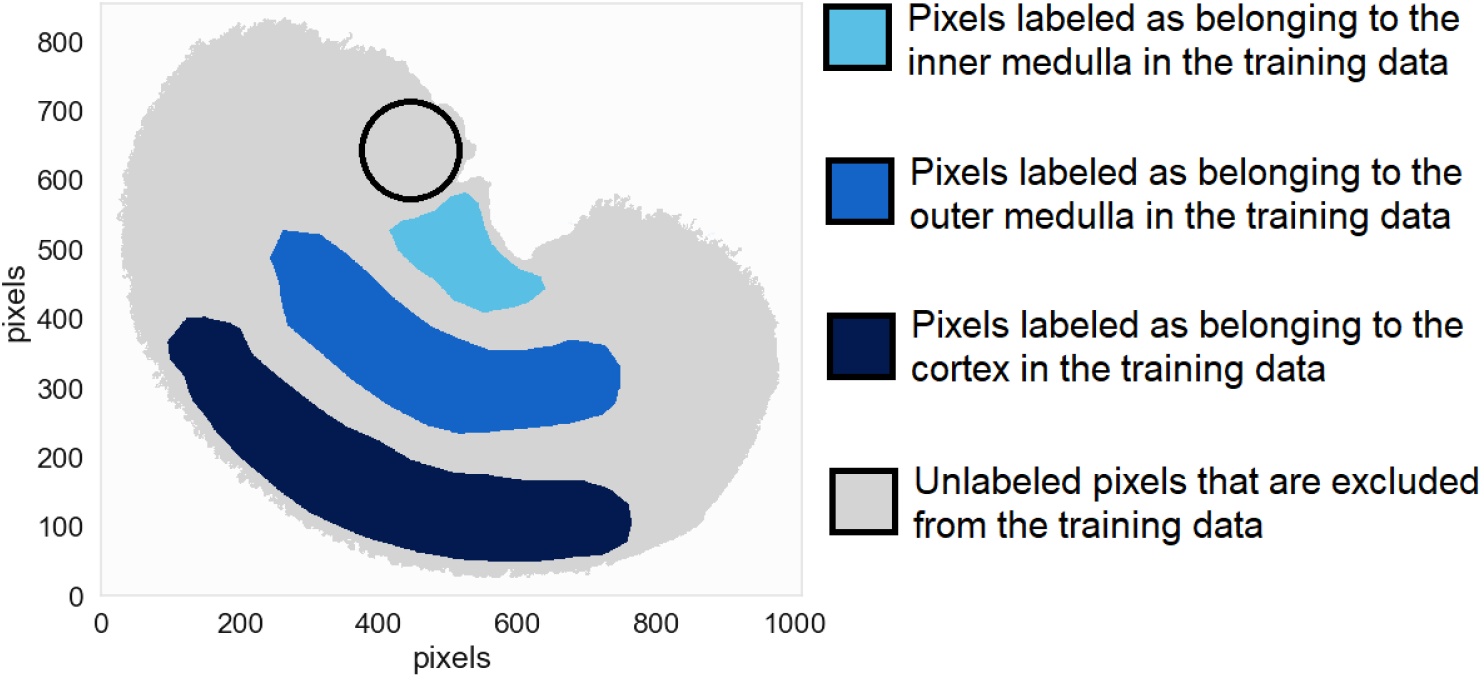
Masks used as inputs for training the three XGBoost classification models directed at recognizing the kidney’s inner medulla, outer medulla, and cortex. Different regions of the tissue sample were manually annotated as belonging to one of four categories: light blue pixels belong to the inner medulla, medium blue pixels belong to the outer medulla, dark blue pixels belong to the cortex, and gray pixels are close to borders between these anatomical structures, making it difficult to annotate them definitively. The latter are excluded from the training data to avoid feeding the model unreliable annotations during training. The black circle outlines a region of the renal cortex that was affected by a sample preparation artefact.

#### Molecular marker discovery for the renal inner medulla

Figure 10a presents the class prediction result for the renal inner medulla model, showing that the inner medulla is successfully differentiated from the outer medulla and cortex. Regarding generalization performance, the XGBoost classification model trained to recognize pixels belonging to the inner medulla achieves a balanced accuracy of 0.9992, a precision of 0.9989, and a recall of 0.9985. Note the slightly noisy region to the top-left of the medulla in Figure 10a. The difficulties encountered by the model in this region, which is outlined by a black circle in Figure 9, are probably due to a sample preparation artefact known as visceral fat delocalization [75]. Figure 10b shows the top ten molecular markers out of the global ranking of 1,428 features (i.e. *m/z* values) as obtained by TreeSHAP. The features are ranked in descending order of global SHAP score, and thus in descending order of relevance to inner medulla tissue recognition. Figure 10b therefore provides insight into a feature’s global relevance to the recognition task of Figure 10a. The most important feature to the XGBoost classification model used to assign pixels to the inner medulla (or not) has a *m/z* value of 1401.043 and a global SHAP score of 2.036.

**Figure 10.**
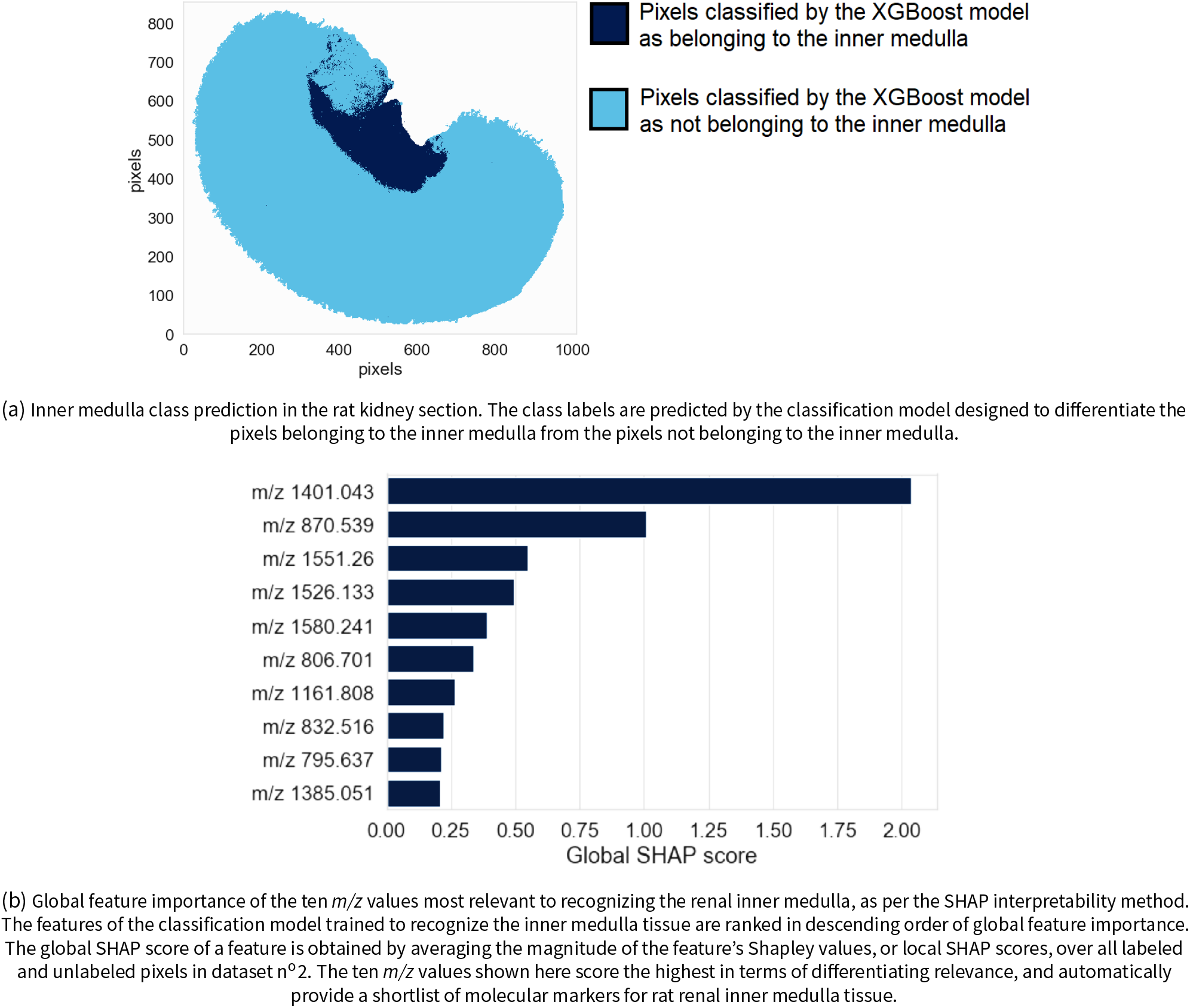
Renal inner medulla recognition and global feature ranking.

The left column of Figure 11 provides information about the spatial distribution and relative abundance of the three top-ranking molecular features for recognizing the inner medulla: Figures 11a, 11c, and 11e are the ion images of the features ranked n°1, °2, and °3 respectively. The right column of Figure 11 provides information on the signs and magnitudes of the local SHAP scores, or Shapley values, of each top-ranking feature across the sample: Figures 11b, 11d, and 11f are the SHAP maps of the three top-ranking features. These SHAP maps provide information on where and how a given ion intensity signal relates to the task of inner medulla tissue recognition. Combining the left and right columns of Figure 11 provides insight into the predictive model’s decision-making process. The signal intensity measured in the inner medulla for features ranked n°1 and n°2 is low (Figures 11a and 11c), and yet their Shapley values are high in the inner medulla (Figures 11b and 11d). These features, *m/z* 1401.043 and *m/z* 870.539 respectively, are negatively correlated to the XGBoost classification model assigning a pixel to the inner medulla. In other words, measuring a low intensity for *m/z* 1401.043 and *m/z* 870.539 in a given pixel increases the log-odds (and probability) of the classification model assigning that pixel to the inner medulla. Given the high predictive performance of the classification model, we can assume that measuring a low intensity for *m/z* 1401.043 and *m/z* 870.539 in a given pixel also increases the probability of that pixel actually belonging to the inner medulla. Conversely, the feature ranked n°3 (*m/z* 1551.260) is positively correlated with the model predicting a pixel as belonging to the inner medulla: its intensities (Figure 11e) and its Shapley values (Figure 11f) are both high in the inner medulla. Measuring a high intensity for *m/z* 1551.260 in a given pixel increases the log-odds (and probability) of the classification model assigning that pixel to the inner medulla. Given the high predictive performance of the classification model, we can assume that measuring a high intensity for *m/z* 1551.260 in a given pixel increases the probability of that pixel actually belonging to the inner medulla. The absence of features ranked n°1 and n°2, and the presence of the feature ranked n°3 seem to be characteristic of renal inner medulla tissue.

**Figure 11.**
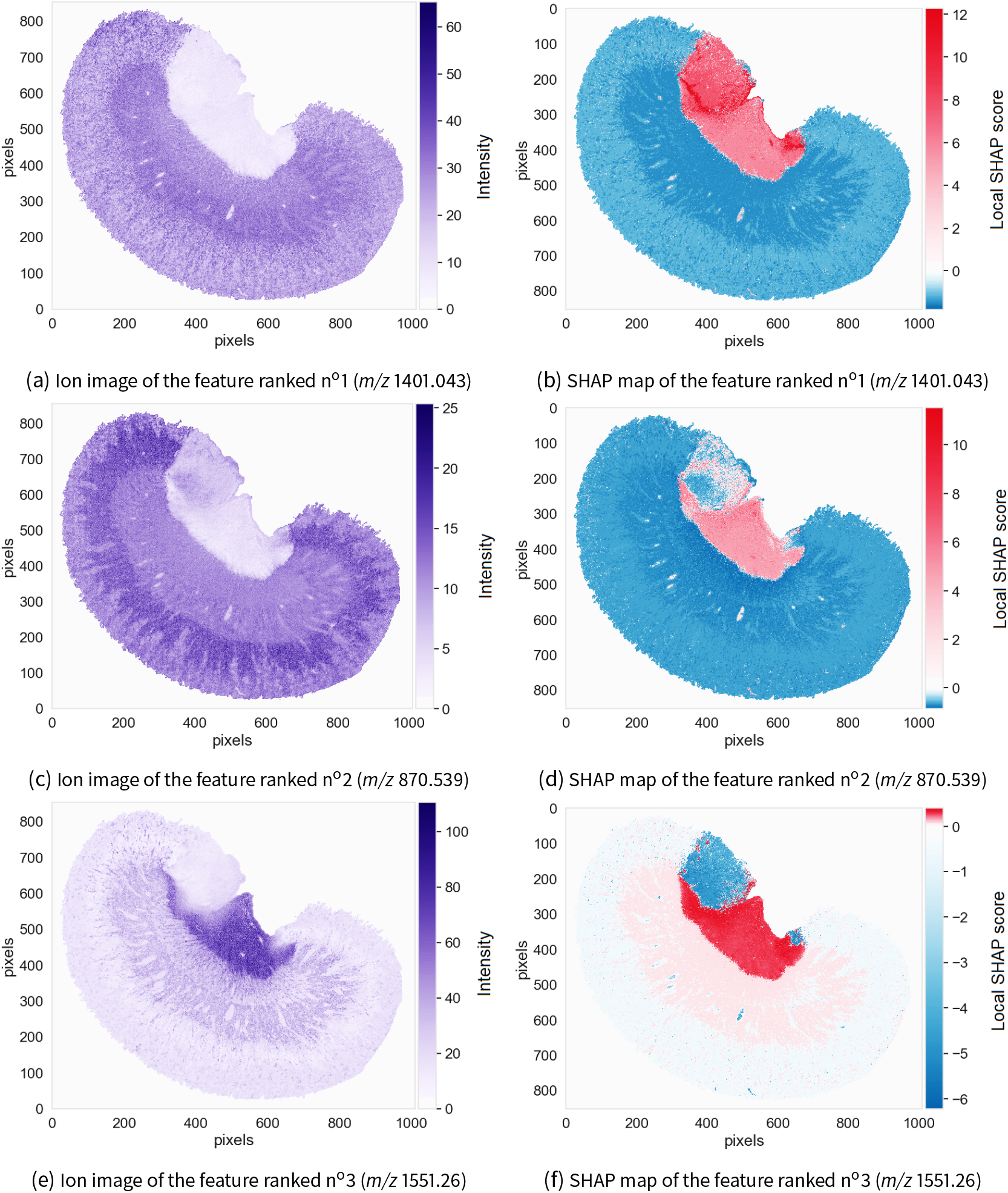
Three promising molecular markers for the renal inner medulla. The ion images (left) and SHAP maps (right) of three features (i.e. *m/z* values) with the most influence on the decision-making process of the classification model trained to recognize the rat’s renal inner medulla are shown. The ion images plot the spatial distribution and measured intensity of each feature across the sample, and are not specifically tied to the task of recognizing the inner medulla. The SHAP maps plot the spatial distribution of Shapley values, or local SHAP predictive importance scores, of each feature across the sample, and provide information on where and how the feature is relevant to the task of recognizing the inner medulla.

We now focus on the tissue region to the top-left of the medulla that actually belongs to the cortex, and that was difficult for the XGBoost classification model to correctly differentiate from the inner medulla (see Figure 10a). The SHAP map of the feature ranked n°1 shows (by coloring the difficult-to-classify area red) that this feature strongly increases the log-odds (and probability) of the cortex pixels to the top-left of the inner medulla being erroneously assigned to the inner medulla: the Shapley values of the feature ranked n°1 are positive with a high magnitude in this region of the cortex. The SHAP map of the feature ranked n°3 shows (by coloring the difficult-to-classify area blue) that the classification model uses this feature to correct for the labeling suggested by the feature ranked n°1: the Shapley values of the feature ranked n°3 in the region to the top-left of the inner medulla are negative with a high magnitude. This case study demonstrates an interesting level of nuance in molecular marker discovery, uniquely provided by the SHAP map representation.

If only the global SHAP scores of the features are taken into account (i.e. only the global information provided in Figure 10b, without the localized information provided in Figures 11b, 11d, and 11f), one might be tempted to consider *m/z* 1401.043 (corresponding to the feature ranked n°1) as the most promising marker candidate for inner medulla tissue in this dataset. Although *m/z* 1401.043 has the most influence on the XGBoost classification model designed to recognize the inner medulla, its global SHAP score is based on a sample-wide assessment of discriminative relevance and disregards subtle spatially localized patterns. In fact, Figure 11b shows that *m/z* 1401.043 has a positive influence on the model’s prediction in the inner medulla but also in a region of the cortex where visceral fat delocalization probably occurred. Unlike *m/z* 1401.043 and *m/z* 870.539 (corresponding to the features ranked n°1 and n°2 respectively), *m/z* 1551.260 (corresponding to feature ranked n°3) is exclusive to the inner medulla. Although the global SHAP scores of *m/z* 1401.043 and *m/z* 870.539 are higher than that of *m/z* 1551.260 (respectively 2.036 and 1.007 versus 0.548), a localized study using SHAP maps shows that *m/z* 1551.260 is the more reliable molecular marker of the three for the renal inner medulla because of its high spatial specificity. Unlike the signal of *m/z* 1551.260, the signals corresponding to *m/z* 1401.043 and *m/z* 870.539 were affected by the sample preparation artefact that took place in the renal cortex. This example also illustrates the importance of not basing one’s estimate of a molecular marker candidate’s relevance exclusively on its global SHAP predictive importance score. When visualized in the form of SHAP maps, the local SHAP scores (or Shapley values) provide useful spatially localized information as to how and where the molecular marker influences the predictive model’s output and (assuming the classification model has good predictive performance) how it ties to the underlying tissue.

## 4 Conclusion

In this work, we propose an innovative computational approach for automating the discovery of biomarker candidates in molecular imaging data. Our approach enables one to efficiently filter a multitude of molecular species down to a panel of promising biomarker candidates. Applying the automated biomarker candidate discovery workflow to imaging mass spectrometry (IMS) data is especially interesting because of the massively multiplexed nature of IMS. By enabling the untargeted concurrent mapping of hundreds to thousands of molecular species across a tissue sample, IMS enables one to cast a wide net for molecular species with biomarker potential. However, the wide range of candidates can pose difficulties since manual examination of IMS data is impractical. Automating biomarker candidate discovery in IMS using machine learning (ML) methodologies, rather than resorting to manual examination, can help re-establish the practical feasibility of IMS-based biomarker discovery, and can help maintain objectivity, scalability, and reproducibility. Our biomarker candidate discovery workflow produces a ranking of molecular species according to the discriminative relevance they hold for a given tissue structure or disease condition, such that the top-ranking molecular species are highly promising biomarker candidates that merit further study.

Our approach to biomarker candidate discovery is to identify highly discriminative molecular species whose overexpression or underexpression characterize a user-defined biological class of interest. A supervised ML algorithm, called extreme gradient boosting (XGBoost), is used to learn a classification model from labeled imaging mass spectrometry data, and a state-of-the-art ML model interpretability method, called Shapley additive explanations (SHAP), is used to measure the local and global predictive importance of the *m/z* values that the model uses as features. We translate the task of biomarker candidate discovery into a feature ranking problem: the features are ranked in descending order of global SHAP importance and the top-ranking features are retained for further investigation. The TreeSHAP implementation of Shapley additive explanations, with observational Shapley values, is used for quantifying the local and global predictive importance of features. In order to add nuance to our analysis, we furthermore introduce SHAP maps, a novel representation and visualization that brings a spatial dimension to our understanding of the decision-making processes of a classification model. The SHAP map of a feature is obtained by plotting that feature’s local SHAP importance scores, or Shapley values, across all pixels making up the sample surface. A feature’s local SHAP importance score is informative of the direction (e.g. positive or negative) and magnitude (e.g. large or small) of the feature’s influence on the classification model’s output for a given pixel. SHAP maps provide insight into the spatial specificity of biomarker candidates by showing how and where a feature influences the classification model’s probability of assigning a pixel, and its corresponding mass spectrum, to the class of interest.

Although our two case studies concern imaging mass spectrometry data, our biomarker candidate discovery workflow is also applicable to other forms of multiplexed imaging data such as multiplexed fluorescence microscopy (e.g. CODEX), imaging mass cytometry, near-infrared imaging, and Raman spectroscopic imaging, and therefore holds the potential to substantially advance biomarker development across a wide range of spectral imaging modalities. One area where our approach can be employed is in the discovery of clinically relevant molecular signatures for functional tissue units in the context of large-scale molecular mapping projects such as the NIH-sponsored Human BioMolecular Atlas Program [76], which aims to build a complete molecular map of the human body at single-cell resolution, and the Kidney Precision Medicine Project [77], which aims to build a comprehensive molecular, cellular, and anatomical map of the kidney. Our work on ML interpretability for multiplexed imaging may also help advance research in biomedical imaging, for example in the field of data-driven multi-modal image fusion [78], where a cross-modal regression model ties the observations in one imaging modality to the observations in another modality. Obtaining spatially-localized insight into how cross-modal connections are made holds potential for advancing all fusion applications, including prediction to a higher spatial resolution, out-of-sample predictions, as well as cross-modal denoising and relationship discovery [79].

## Nomenclature

IMS: imaging mass spectrometry
*m/z*: mass-to-charge ratio
MALDI: matrix assisted laser desorption/ionization
ML: machine learning
PI: permutation importance
Q-TOF: quadrupole time-of-flight
SHAP: Shapley additive explanations
XGBoost: extreme gradient boosting

## Acknowledgements

The authors thank Elizabeth Neumann, Kavya Sharman, Mark deCaestecker, Roger Moens, and Emilio Rivera for their support and helpful suggestions. Research reported in this publication was supported by the National Institutes of Health (NIH)’s Common Fund, National Institute Of Diabetes And Digestive And Kidney Diseases (NIDDK), and the Office Of The Director (OD) under Award Number U54DK120058 (J.M.S., R.M.C., and R.V.), by NIH’s Common Fund, National Eye Institute, and the Office Of The Director (OD) under Award Number U54EY032442 (J.M.S., R.M.C., and R.V.), by NIH’s National Institute Of Allergy And Infectious Diseases (NIAID) under Award Numbers R01AI138581 and R01AI145992 (J.M.S. and R.V.), by NIH’s National Institute of General Medical Sciences (NIGMS) under Award Number P41GM103391 (R.M.C.), and by the National Science Foundation Major Research Instrument Program CBET – 1828299 (J.M.S. and R.M.C.). The research was furthermore supported by the Nederlandse Organisatie voor Wetenschappelijk Onderzoek (NWO; Dutch Research Council), ZonMw, FLAG-ERA, and the European Commission through the FLAG-ERA III JTC project SMART BRAIN (NWO 680-91-319) under the NWO-domain Exacte en Natuurwetenschappen, together with the NWO-domain Sociale en Geesteswetenschappen, and in association with the European Commission’s Human Brain Project (R.V.). The content is solely the responsibility of the authors and does not necessarily represent the official views of the National Institutes of Health, the Nederlandse Organisatie voor Wetenschappelijk Onderzoek, ZonMw, FLAG-ERA, or the European Commission.

## Supplementary material

Additional information for the paper entitled “Automated Biomarker Candidate Discovery in Imaging Mass Spectrometry Data Through Spatially Localized Shapley Additive Explanations” by Leonoor E.M. Tideman, Lukasz G. Migas, Katerina V. Djambazova, Nathan Heath Patterson, Richard M. Caprioli, Jeffrey M. Spraggins, and Raf Van de Plas. Corresponding author

Contents:

1. Experimental protocol
2. Data preprocessing
3. Additional case studies

### Experimental protocol

#### Materials

Acetic acid, 1,5-diaminonaphthalene (DAN), ammonium formate, hematoxylin, and eiosin were purchased from Sigma-Aldrich Chemical Co. (St. Louis, MO, USA). HPLC-grade ethanol was purchased from Fisher Scientific (Pittsburgh, PA, USA).

#### Sample Preparation

One-week old C57BL/6 control mouse pup was snap frozen at −80oC, shaved over dry ice, and cryosectioned at 20 μm thickness, using a CM3050 S cryostat (Leica Biosystems, Wetzlar, Germany). The tissue was thaw-mounted onto a conductive indium tin oxide coated glass slide (Delta Technologies, Loveland, CO, USA). Rat kidney tissue was purchased from PelFreeze Biologicals (Rogers, AR, USA), sectioned at 10 μm thickness and thaw-mounted onto a conductive slide. Autofluorescence microscopy images were acquired using EGFP, DAPI, and DsRed filters on a Zeiss AxioScan Z1 slide scanner (Carl Zeiss Microscopy GmbH, Oberkochen, Germany). Approximately 500 mg of DAN was sublimed at 130oC and 24 mTorr for 3.5 min onto the tissue surface for a final density of ~1.0 mg/cm^2^.

#### MALDI TIMS-IMS

Our biomarker candidate discovery workflow is demonstrated on two imaging mass spectrometry (IMS) datasets. All experiments were carried out on a prototype timsTOF fleX mass spectrometer (Bruker Daltonik, Bremen, Germany).

- Dataset n°1: Mouse pup images were acquired in trapped ion mobility (TIMS) mode of operation with an ion transfer time of 100 μs, prepulse storage time of 8 μs, and a collision RF of 2,000 Vpp, a TIMS funnel 1 (accumulation) RF of 450 Vpp, a TIMS funnel 2 RF (analysis) of 400 Vpp, a multipole RF of 400 Vpp, and a collision cell entrance (in) voltage of 300 V. Tissue imaging data (164,808 pixels) were collected at 50 μm spatial resolution, using 200 shots per pixel and 48% laser power. Data were collected in positive ionization mode from *m/z* 300 to 1,200. The TIMS scan time was set to 400 ms, with a reduced mobility (1/K_0_) range of 0.4 - 1.9 (V·s)/cm2.
- Dataset n°2: Murine kidney images were generated in Q-TOF mode of operation. Tissue imaging data (591,534 pixels) were collected at 15 μm spatial resolution, using 400 shots per pixel, and 35% laser power. Data were collected in positive ionization mode from *m/z* 200 to 1,500.

#### Histology

Following MALDI IMS, matrix was removed from tissue using 100% ethanol and rehydrated with graded ethanol and double distilled H2O. The tissues were then stained using a hematoxylin and eosin (H&E) stain. Brightfield microscopy of stained tissues was obtained at 20× magnification using a Leica SCN400 Brightfield Slide Scanner (Leica Microsystems, Buffalo Grove, IL, USA).

### Data preprocessing

- Dataset n°1: The mouse-pup data was exported into a custom binary format optimized for storage and speed of analysis of the ion mobility-IMS data. Each frame/pixel contains between 10,000-100,000 centroid peaks that span the acquisition range of *m/z* 300-1,200 and 1/K_0_ 0.4-1.9 (V·s)/cm2 with 221,888 and 4,001 bins in the mass spectrometry and ion mobility dimensions, respectively. The processing pipeline requires common *m/z* and 1/K_0_ axes, hence individual centroid peaks were inserted at their correct bin positions along the mass spectrometry and ion mobility dimensions; missing values were set to zero. Following the conversion process, a mean mass spectrum of the entire dataset was generated, and peak picked. A total of 879 features were selected and extracted to generate ion mobility-rich ion images. Since our biomarker candidate discovery workflow is optimized for non-ion mobility data, the ion mobility information is removed by summation of all ion mobility bins of the ion mobility-rich images to a single vector, resulting in a standard ion image.
- Dataset n°2: The murine kidney data was exported into a custom binary format as described above. Data were acquired in the Q-TOF only mode, hence the ion mobility dimension is not present, in which case we introduce secondary dimension by enforcing the dataset to contain one ion mobility bin. This is carried out to ensure efficient storage and data processing without having any impact on the actual data. Following the conversion process, a mean mass spectrum of the entire dataset was generated, and peak picked resulting in a total of 1,428 ion images.

Low variance noise was removed from both datasets by principal component analysis: principal component analysis is a matrix factorization technique that can be used to reduce the dimensionality of an IMS dataset while retaining most of its original variance, and hence most of its latent molecular information [21, 80]. Feature centering, which consists in subtracting the mean from each column of the IMS data matrix, is performed without feature scaling. We argue that feature scaling prior to classification model training is not necessary because large differences in high intensity peaks are biologically informative, and high intensity peaks should therefore have more influence on the model than low intensity peaks [21].

### Additional case studies: IMS dataset n°2

#### Biomarker candidate discovery for the outer renal medulla

Figure S1a presents the classification result: the outer medulla has successfully been differentiated from the cortex and inner medulla. Regarding generalization performance, the XGBoost classification model trained to recognize pixels belonging to the outer renal medulla achieves a balanced accuracy of 0.9970, a precision of 0.9972 and a recall of 0.9958.

**Figure S1.**
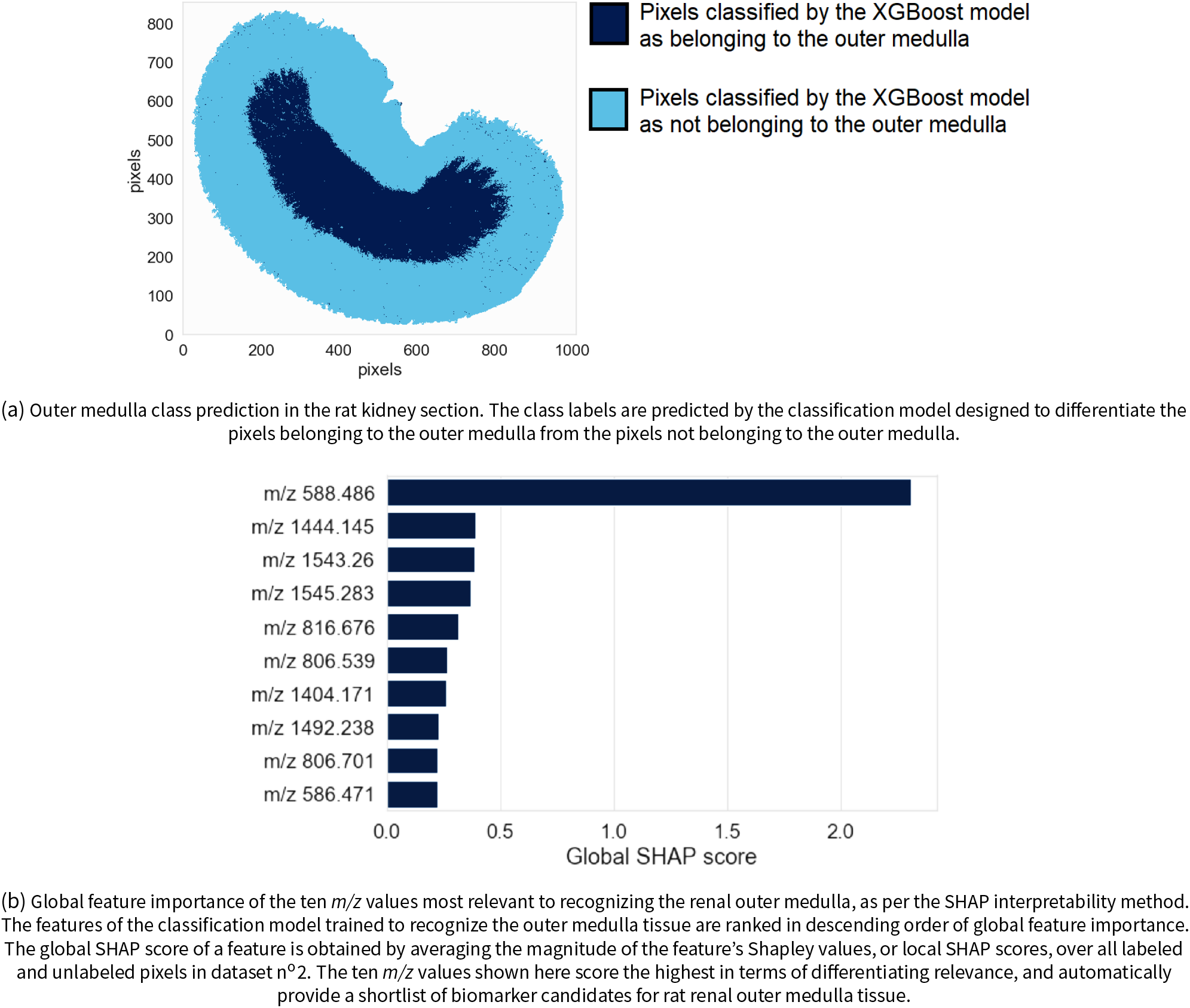
Renal outer medulla recognition and global feature ranking.

Figure S1b shows the top ten molecular markers out of the global ranking of 1428 features (i.e. *m/z* values) as obtained by TreeSHAP. The features are ranked in descending order of global SHAP score, and thus in descending order of relevance to outer medulla tissue recognition. The model used to assign pixels to the outer medulla (or not) relies heavily on the feature ranked n°1, whose *m/z* value is 588.486 and whose global SHAP score of 2.311. The left column of Figure S2 provides information about the spatial distribution and relative abundance of the three top-ranking molecular features for recognizing the outer medulla. The right column of Figure S2 provides information on the signs and magnitudes of the local SHAP scores, or Shapley values, of each top-ranking feature across the sample. These SHAP maps provide information on where and how a given ion intensity signal relates to the task of outer medulla tissue recognition. The feature ranked n°1 is strongly positively correlated with assigning a pixel to the outer medulla because its ion image (Figure S2a) shows a high intensity in the outer medulla and because its SHAP map (Figure S2b) shows positive Shapley values in the outer medulla and negative Shapley values in the inner medulla and cortex. Given the high predictive performance of the classification model, we can assume that measuring a high intensity of the feature ranked n°1 in a given pixel also increases the probability of that pixel actually belonging to the outer medulla. Features ranked n°2 and n°3 (*m/z* 1444.145 and *m/z* 1543.26 respectively) are negatively correlated with the model assigning a pixel to the outer medulla. Given the high predictive performance of the classification model, we can assume that measuring a low intensity of the features ranked n°2 and n°3 in a given pixel also increases the probability of that pixel actually belonging to the outer medulla. The SHAP maps of features ranked n°2 and n°3 (Figures S2d and S2f respectively) show that the XGBoost classification model uses the feature ranked n°2 to differentiate the outer medulla from the inner medulla and cortex, and that it uses the feature ranked n°3 to differentiate the outer medulla and inner medulla from the cortex.

**Figure S2.**
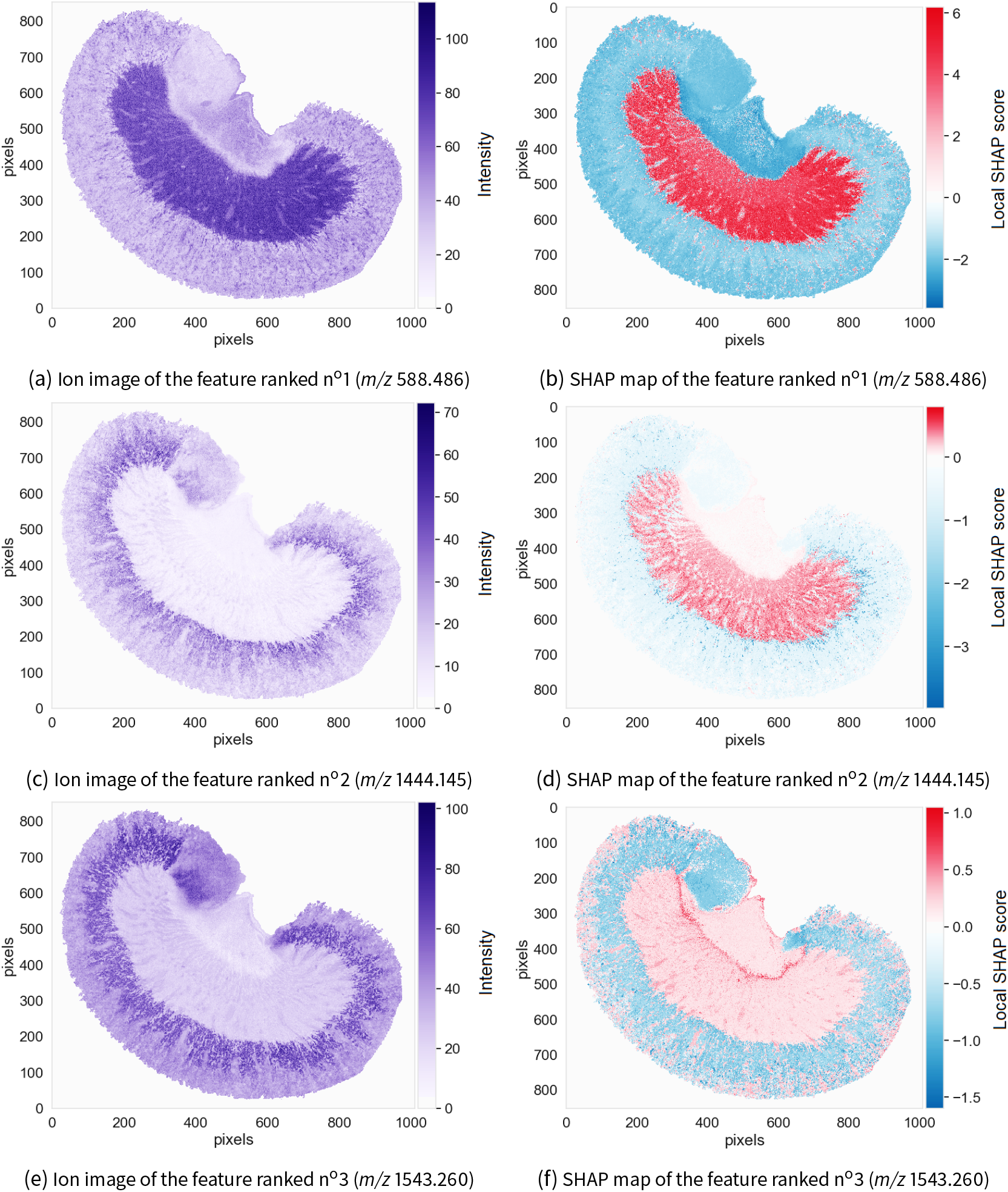
Three promising molecular markers for the renal outer medulla. The ion images (left) and SHAP maps (right) of three features (i.e. *m/z* values) with the most influence on the decision-making process of the classification model trained to recognize the rat’s renal outer medulla are shown. The ion images plot the spatial distribution and measured intensity of each feature across the sample, and are not specifically tied to the task of recognizing the outer medulla. The SHAP maps plot the spatial distribution of Shapley values, or local SHAP predictive importance scores, of each feature across the sample, and provide information on where and how the feature is relevant to the task of recognizing outer medulla.

#### Biomarker candidate discovery for the renal cortex

Figure S3a presents the classification result. The cortex has successfully been differentiated from the inner and outer medulla. Regarding generalization performance, the XGBoost classification model trained to recognize pixels belonging to the renal cortex achieves a balanced accuracy of 0.9974, a precision of 0.9976 and a recall of 0.9973.

**Figure S3.**
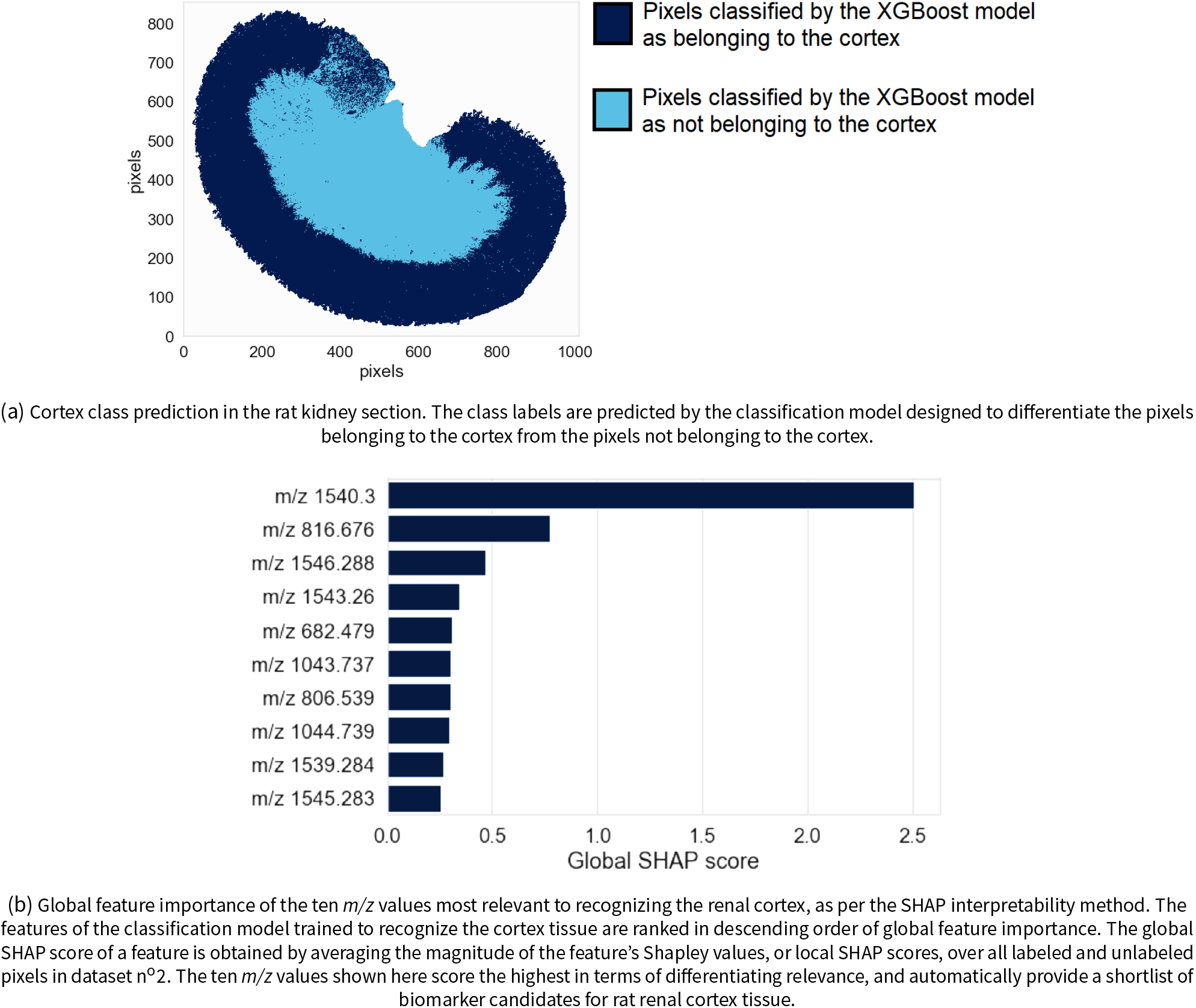
Renal cortex recognition and global feature ranking.

Figure S3b shows the top ten molecular markers out of the global ranking of 1428 features (i.e. *m/z* values) as obtained by TreeSHAP. The features are ranked in descending order of global SHAP score, and thus in descending order of relevance to cortex tissue recognition. The XGBoost model used to assign pixels to the cortex (or not) relies heavily upon one feature, whose *m/z* value is 1540.300 and whose global SHAP score is 2.507. The left column of Figure S4 presents the ion images of the three top-ranking features. The right column of Figure S4 presents the SHAP maps of the three top-ranking features. The ion image of the feature ranked n°1 (Figure S4a) shows that feature ranked n°1 has high intensity in the cortex; and the SHAP map of feature ranked n°1 (Figure S4b) shows that feature ranked n°1 has positive Shapley values in the cortex, and negative Shapley values in the inner and outer medulla. Therefore, feature ranked n°1 (i.e. *m/z* 1540.300) is positively correlated to a pixel being assigned to the cortex. Conversely, features ranked n°2 and n°3 (i.e. *m/z* 816.676 and *m/z* 1546.288 respectively) are negatively correlated with the XGBoost classification model assigning a pixel to the cortex. Given the high predictive performance of the classification model, we can assume that measuring a high intensity of feature ranked n°1, and a low intensity of features ranked n°2 and n°3, increase the probability of that pixel actually belonging to the cortex.

**Figure S4.**
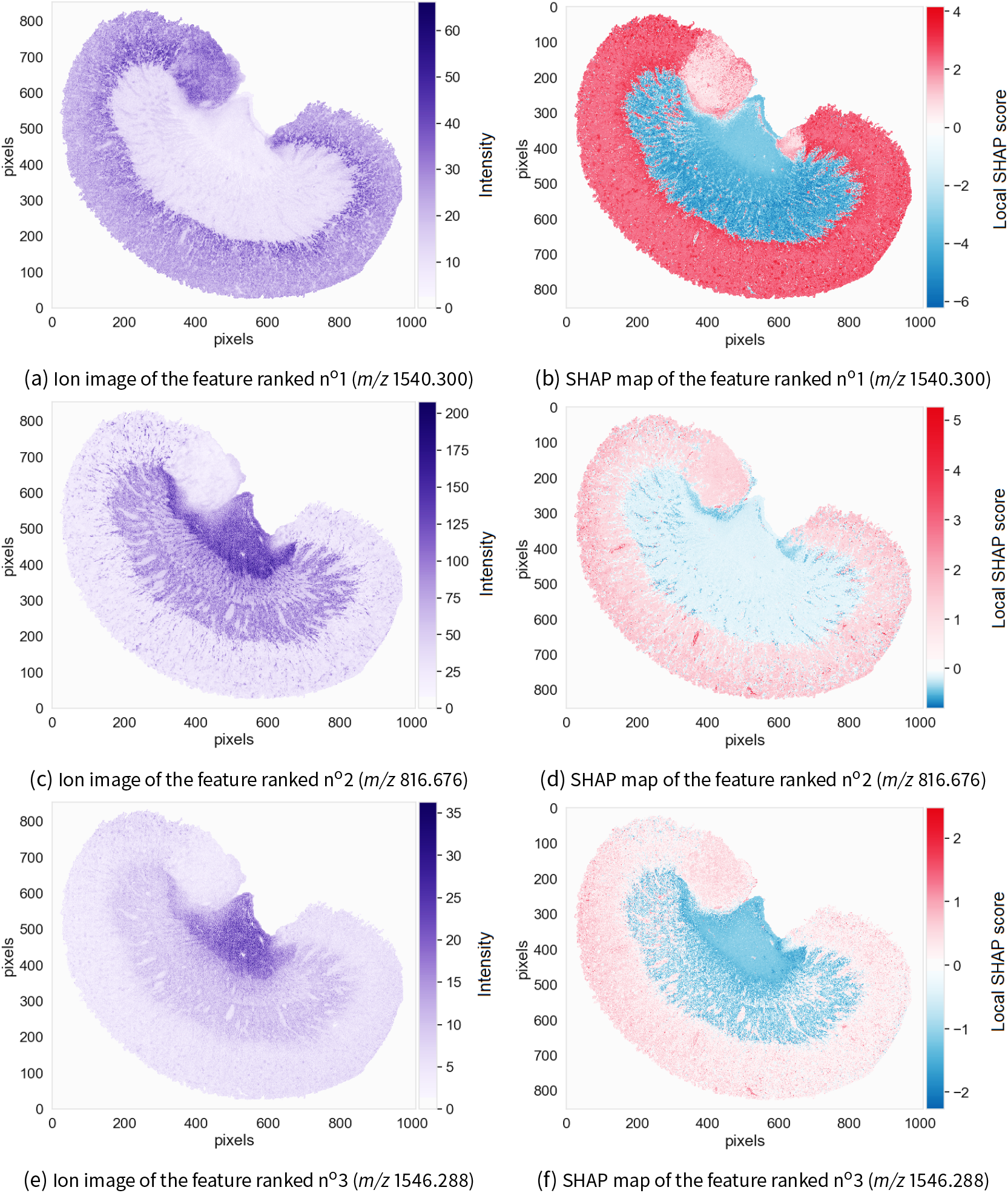
Three promising molecular markers for the renal cortex. The ion images (left) and SHAP maps (right) of three features (i.e. *m/z* values) with the most influence on the decision-making process of the classification model trained to recognize the rat’s renal cortex are shown. The ion images plot the spatial distribution and measured intensity of each feature across the sample, and are not specifically tied to the task of recognizing the cortex. The SHAP maps plot the spatial distribution of Shapley values, or local SHAP predictive importance scores, of each feature across the sample, and provide information on where and how the feature is relevant to the task of recognizing the cortex.

## Footnotes

1 There is no formal definition of supervised machine learning model interpretability that is agreed-upon within the computer science and data science communities [26, 27, 28]. The definition we propose, namely explaining a model’s decision-making process by quantifying the influence of its input features on its output, is specific to the analysis of imaging mass spectrometry data using supervised machine learning methods other than deep learning. We use the terms interpretability and explainability interchangeably.

2 Overfitting is a common issue that occurs when a supervised machine learning model adapts too closely to the training data and memorizes not only the relationship between inputs and outputs but also the noise. The structure of an overfit model is very sensitive to changes in its training data. Such a model usually performs poorly on new data. The risk of overfitting tends to increase when handling high-dimensional data like imaging mass spectrometry data.

3 Gini importance, which is the default measure of feature importance in Scikit-Learn’s implementation of random forest [64], and gain, which is the default measure of feature importance in the Scikit-Learn wrapper interface for XGBoost’s implementation of extreme gradient boosting [65], are two examples of popular yet inconsistent impurity-based approaches for estimating features’ predictive importances. The Gini importance of a feature is computed by averaging the weighted decrease in node impurity achieved by splitting a node using that feature over all decision trees making up the ensemble [66].

4 Permutation importance was originally developed by Breiman, under the name of mean decrease accuracy, as a model-specific method for measuring feature importance in random forests [68]. The idea is to randomly permute a feature across all observations to break its association with the model prediction (and the other features) and effectively cancel its predictive power [69]. Therefore, if the feature under study is strongly associated to the prediction, permuting its values should result in a large drop in predictive performance. Conversely, if the feature is weakly associated to the prediction, permuting it should have little to no impact on performance.

## References

[1] B. D. W. Group. “Biomarkers and Surrogate Endpoints: Preferred Definitions and Conceptual Framework”. en. In: Clinical Pharmacology & Therapeutics 69.3 (2001), pp. 89–95. issn: 1532-6535. doi: 10.1067/mcp.2001.113989.

[2] N. Rifai, M. A. Gillette, and S. A. Carr. “Protein Biomarker Discovery and Validation: The Long and Uncertain Path to Clinical Utility”. In: Nature Biotechnology 24.8 (Aug. 2006), pp. 971–983. issn: 1546-1696. doi: 10.1038/nbt1235.

[3] C. A. Crutchfield et al. “Advances in Mass Spectrometry-Based Clinical Biomarker Discovery”. In: Clinical Proteomics 13 (Jan. 2016). issn: 1542-6416. doi: 10.1186/s12014-015-9102-9.

[4] Z.-Z. Hu et al. “Omics-Based Molecular Target and Biomarker Identification”. In: Methods in molecular biology (Clifton, N.J.) 719 (2011), pp. 547–571. issn: 1064-3745. doi: 10.1007/978-1-61779-027-0_26.

[5] M. Dilillo, B. Heijs, and L. A. McDonnell. “Mass Spectrometry Imaging: How Will It Affect Clinical Research in the Future?” In: Expert Review of Proteomics 15.9 (Sept. 2018), pp. 709–716. issn: 1478-9450. doi: 10.1080/14789450.2018.1521278.

[6] K. Yanagisawa et al. “Proteomic Patterns of Tumour Subsets in Non-Small-Cell Lung Cancer”. In: 362.9382 (Aug. 2003), pp. 433–439. issn: 0140-6736, 1474-547X. doi: 10.1016/S0140-6736(03)14068-8.

[7] R. M. Caprioli, T. B. Farmer, and J. Gile. “Molecular Imaging of Biological Samples: Localization of Peptides and Proteins Using MALDI-TOF MS”. In: Analytical Chemistry 69.23 (Dec. 1997), pp. 4751–4760. issn: 0003-2700. doi: 10.1021/ac970888i.

[8] K. Schwamborn and R. M. Caprioli. “MALDI Imaging Mass Spectrometry – Painting Molecular Pictures”. In: Molecular Oncology. Thematic Issue: Oncoproteomics 4.6 (Dec. 2010), pp. 529–538. issn: 1574-7891. doi: 10.1016/j.molonc.2010.09.002.

[9] S. S. Rubakhin et al. “Imaging Mass Spectrometry: Fundamentals and Applications to Drug Discovery.” In: Drug discovery today 10.12 (2005), pp. 823–837. doi: 10.1016/s1359-6446(05)03458-6.

[10] K. Schwamborn, M. Kriegsmann, and W. Weichert. “MALDI Imaging Mass Spectrometry - From Bench to Bedside”. eng. In: Biochimica Et Biophysica Acta. Proteins and Proteomics 1865.7 (July 2017), pp. 776–783. issn: 1570-9639. doi: 10.1016/j.bbapap.2016.10.014.

[11] M. Stoeckli et al. “Imaging Mass Spectrometry: A New Technology for the Analysis of Protein Expression in Mammalian Tissues”. en. In: Nature Medicine 7.4 (Apr. 2001), pp. 493–496. issn: 1546-170X. doi: 10.1038/86573.

[12] T. Alexandrov. “MALDI Imaging Mass Spectrometry: Statistical Data Analysis and Current Computational Challenges”. In: BMC Bioinformatics 13.Suppl 16 (Nov. 2012), S11. issn: 1471-2105. doi: 10.1186/1471-2105-13-S16-S11.

[13] T. A. Zimmerman et al. “Imaging of Cells and Tissues with Mass Spectrometry: Adding Chemical Information to Imaging”. In: Methods in cell biology 89 (2008), pp. 361–390. issn: 0091-679X. doi: 10.1016/S0091-679X(08)00613-4.

[14] M. Aichler and A. Walch. “MALDI Imaging Mass Spectrometry: Current Frontiers and Perspectives in Pathology Research and Practice”. In: Laboratory Investigation; a Journal of Technical Methods and Pathology 95.4 (Apr. 2015), pp. 422–431. issn: 1530-0307. doi: 10.1038/labinvest.2014.156.

[15] K. Schwamborn. “Imaging Mass Spectrometry in Biomarker Discovery and Validation”. In: Journal of Proteomics 75.16 (Aug. 2012), pp. 4990–4998. issn: 1876-7737. doi: 10.1016/j.jprot.2012.06.015.

[16] S. Lou et al. “High-grade Sarcoma Diagnosis and Prognosis: Biomarker Discovery by Mass Spectrometry Imaging”. In: Proteomics 16 (May 2016), pp. 1802–1813. doi: 10.1002/pmic.201500514.

[17] R. Lazova et al. “Imaging Mass Spectrometry – a New and Promising Method to Differentiate Spitz Nevi from Spitzoid Malignant Melanomas”. In: The American Journal of Dermatopathology 34.1 (Feb. 2012), pp. 82–90. issn: 0193-1091. doi: 10.1097/DAD.0b013e31823df1e2.

[18] L. S. Eberlin et al. “Classifying Human Brain Tumors by Lipid Imaging with Mass Spectrometry”. In: Cancer Research (Dec. 2011). issn: 0008-5472, 1538-7445. doi: 10.1158/0008-5472.CAN-11-2465.

[19] T. Li et al. “In Situ Biomarker Discovery and Label-Free Molecular Histopathological Diagnosis of Lung Cancer by Ambient Mass Spectrometry Imaging”. In: Nature Scientific Reports 5 (Sept. 2015). doi: 10.1038/srep14089.

[20] E. Robotti, M. Manfredi, and E. Marengo. “Biomarkers Discovery through Multivariate Statistical Methods: A Review of Recently Developed Methods and Applications in Proteomics”. In: Journal of Proteomics & Bioinformatics 0.0 (2015), pp. 1–20. issn: 0974-276X. doi: 10.4172/jpb.S3-003.

[21] N. Verbeeck, R. M. Caprioli, and R. Van de Plas. “Unsupervised Machine Learning for Exploratory Data Analysis in Imaging Mass Spectrometry”. In: Mass Spectrometry Reviews (Oct. 2019). doi: 10.1002/mas.21602.

[22] M. Hanselmann et al. “Toward Digital Staining Using Imaging Mass Spectrometry and Random Forests”. In: Journal of proteome research 8.7 (July 2009), pp. 3558–3567. issn: 1535-3893. doi: 10.1021/pr900253y.

[23] C. Molnar, G. Casalicchio, and B. Bischl. “Interpretable Machine Learning – A Brief History, State-of-the-Art and Challenges”. In: arXiv:2070.09337 [cs, stat] (Oct. 2020). arXiv: 2010.09337 [cs, stat].

[24] V. Belle and I. Papantonis. “Principles and Practice of Explainable Machine Learning”. In: arXiv:2009.77698 [cs, stat] (Sept. 2020). arXiv: 2009.11698 [cs, stat].

[25] M. Du, N. Liu, and X. Hu. “Techniques for Interpretable Machine Learning”. In: arXiv:7808.00033 [cs, stat] (May 2019). arXiv: 1808.00033 [cs, stat].

[26] Z. C. Lipton. “The Mythos of Model Interpretability”. In: arXiv:7606.03490 [cs.LG] (Mar. 2017).

[27] F. Doshi-Velez and B. Kim. “Towards A Rigorous Science of Interpretable Machine Learning”. In: arXiv:7702.08608 [cs, stat] (Feb. 2017). arXiv: 1702.08608 [cs, stat].

[28] W. J. Murdoch et al. “Interpretable Machine Learning: Definitions, Methods, and Applications”. In: Proceedings of the National Academy of Sciences 116.44 (Oct. 2019), pp. 22071–22080. issn: 0027-8424, 1091-6490. doi: 10.1073/pnas.1900654116. arXiv: 1901.04592.

[29] C. Molnar. “Chapter 2 Interpretability”. In: Interpretable Machine Learning: A Guide for Making Black Box Models Explainable. Online book: https://christophm.github.io/interpretable-ml-book/, 2020.

[30] C. B. Azodi, J. Tang, and S.-H. Shiu. “Opening the Black Box: Interpretable Machine Learning for Geneticists”. In: Trends in Genetics 36.6 (June 2020), pp. 442–455. issn: 0168-9525. doi: 10.1016/j.tig.2020.03.005.

[31] Y. R. Xie et al. “Single-Cell Classification Using Mass Spectrometry Through Interpretable Machine Learning”. In: Analytical Chemistry (June 2020). issn: 0003-2700. doi: 10.1021/acs.analchem.0c01660.

[32] BioRender. https://biorender.com/.

[33] M. Tharapornsakulwong and Angela. Noun Project: Machine Learning & Mechanism Icons. https://thenounproject.com/.

[34] S. Meding et al. “Tumor Classification of Six Common Cancer Types Based on Proteomic Profiling by MALDI Imaging”. In: Journal of Proteome Research 11.3 (Mar. 2012), pp. 1996–2003. issn: 1535-3907. doi: 10.1021/pr200784p.

[35] Y. Zhang and X. Liu. “Machine Learning Techniques for Mass Spectrometry Imaging Data Analysis and Applications”. In: Bioanalysis 10.8 (Mar. 2018), pp. 519–522. issn: 1757-6180. doi: 10.4155/bio-2017-0281.

[36] M. Galli et al. “A Support Vector Machine Classification of Thyroid Bioptic Specimens Using MALDI-MSI Data”. In: 2016 (2016). issn: 1687-8027. doi: 10.1155/2016/3791214.

[37] J. Behrmann et al. “Deep Learning for Tumor Classification in Imaging Mass Spectrometry”. In: Bioinformatics (Oxford, England) 34.7 (Jan. 2018), pp. 1215–1223. issn: 1367-4811.

[38] J. van Kersbergen et al. “Cancer Detection in Mass Spectrometry Imaging Data by Dilated Convolutional Neural Networks”. In: Medical Imaging 2079: Digital Pathology. Vol. 10956. International Society for Optics and Photonics, Mar. 2019, p. 109560I. doi: 10.1117/12.2512360.

[39] Z. Zhou and R. N. Zare. “Personal Information from Latent Fingerprints Using Desorption Electrospray Ionization Mass Spectrometry and Machine Learning”. In: Analytical Chemistry 89.2 (Jan. 2017), pp. 1369–1372. issn: 1520-6882. doi: 10.1021/acs.analchem.6b04498.

[40] K. Margulis et al. “Combining Desorption Electrospray Ionization Mass Spectrometry Imaging and Machine Learning for Molecular Recognition of Myocardial Infarction”. In: Analytical Chemistry 90.20 (Oct. 2018), pp. 12198–12206. issn: 1520-6882. doi: 10.1021/acs.analchem.8b03410.

[41] M. Krzywinski and N. Altman. “Classification and Regression Trees”. en. In: Nature Methods 14.8 (Aug. 2017), pp. 757–758. issn: 1548-7105. doi: 10.1038/nmeth.4370.

[42] R. O. Duda, D. G. Stork, and P. E. Hart. “Chapter 8: Tree-Based Methods”. In: Pattern Classification. Second. John Wiley & Sons, 2012.

[43] L. Breiman et al. Classification and Regression Trees. Taylor & Francis, 1984. isbn: 978-0-412-04841-8.

[44] S. Russell and P. Norvig. “Chapter 18: Learning from Examples”. In: Artificial Intelligence, a Modern Approach. 3rd. Prentice Hall. isbn: 978-0-13-604259-4.

[45] T. Chen and C. Guestrin. “XGBoost: A Scalable Tree Boosting System”. In: Proceedings of the 22nd ACM SIGKDD International Conference on Knowledge Discovery and Data Mining - KDD ‘76 (2016), pp. 785–794. doi: 10.1145/2939672.2939785. arXiv: 1603.02754.

[46] J. Friedman. “Stochastic Gradient Boosting”. In: Computational Statistics & Data Analysis 38 (Mar. 1999), pp. 367–378. doi: 10.1016/S0167-9473(01)00065-2.

[47] J. H. Friedman. “Greedy Function Approximation: A Gradient Boosting Machine.” In: Annals of Statistics 29.5 (Oct. 2001), pp. 1189–1232. issn: 0090-5364, 2168-8966. doi: 10.1214/aos/1013203451.

[48] Y. Freund and R. E. Schapire. “A Short Introduction to Boosting”. In: In Proceedings of the Sixteenth International Joint Conference on Artificial Intelligence. Morgan Kaufmann, 1999, pp. 1401–1406.

[49] K. P. Murphy. “Chapter 16: Adaptive Basis Function Models (Section 16.4: Boosting & Section 16.3: Generalized Additive Models)”. In: Machine Learning: A Probabilistic Perspective. The MIT Press, 2012. isbn: 978-0-262-01802-9.

[50] J. Friedman, T. Hastie, and R. Tibshirani. “Additive Logistic Regression: A Statistical View of Boosting”. In: The Annals of Statistics 28 (Apr. 2000), pp. 337–407. doi: 10.1214/aos/1016218223.

[51] T. Hastie, R. Tibshirani, and J. Friedman. “Chapter 10: Boosting and Additive Trees”. In: Elements of Statistical Learning: Data Mining, Inference, and Prediction. Second. Springer-Verlag, 2017. isbn: 978-0-387-84857-0.

[52] T. Hastie, R. Tibshirani, and J. Friedman. “Chapter 9: Additive Models, Trees, and Related Methods”. In: Elements of Statistical Learning: Data Mining, Inference, and Prediction. Second. Springer-Verlag, 2017. isbn: 978-0-387-84857-0.

[53] M. Kuhn and K. Johnson. “Chapter 8: Regression Trees and Rule-Based Models (Section 8.6: Boosting)”. In: Applied Predictive Modeling. 1st. Springer, 2013. isbn: ISBN 978-1-4614-6849-3.

[54] XGBoost Parameters — Xgboost 7.2.0-SNAPSHOT Documentation. https://xgboost.readthedocs.io/en/latest/parameter.html.

[55] Python API Reference — Xgboost 7.2.0-SNAPSHOT Documentation. https://xgboost.readthedocs.io/en/latest/python/python_api.html#module-xgboost.core.

[56] S. Lundberg and S.-I. Lee. “An Unexpected Unity among Methods for Interpreting Model Predictions”. In: arXiv:7677.07478 [cs] (Nov. 2016). arXiv: 1611.07478 [cs].

[57] S. Lundberg and S.-I. Lee. “A Unified Approach to Interpreting Model Predictions”. In: arXiv:7705.07874 [cs, stat] (May 2017). arXiv: 1705.07874 [cs, stat].

[58] E. Strumbelj and I. Kononenko. “An Efficient Explanation of Individual Classifications Using Game Theory”. In: J. Mach. Learn. Res. 11 (Mar. 2010), pp. 1–18. issn: 1532-4435.

[59] E. Strumbelj and I. Kononenko. “Explaining Prediction Models and Individual Predictions with Feature Contributions”. In: 41.3 (Dec. 2014), pp. 647–665. issn: 0219-3116. doi: 10.1007/s10115-013-0679-x.

[60] M. T. Ribeiro, S. Singh, and C. Guestrin. “Why Should I Trust You?: Explaining the Predictions of Any Classifier”. In: Proceedings of the 22nd ACM SIGKDD International Conference on Knowledge Discovery and Data Mining - KDD ‘76. San Francisco, California, USA: ACM Press, 2016, pp. 1135–1144. isbn: 978-1-4503-4232-2. doi: 10.1145/2939672.2939778.

[61] S. M. Lundberg et al. “From Local Explanations to Global Understanding with Explainable AI for Trees”. In: Nature Machine Intelligence 2.1 (Jan. 2020), pp. 56–67. issn: 2522-5839. doi: 10.1038/s42256-019-0138-9.

[62] H. Chen et al. “True to the Model or True to the Data?” In: arXiv:2006.76234 [cs, stat] (June 2020). arXiv: 2006.16234 [cs, stat].

[63] S. M. Lundberg, G. G. Erion, and S.-I. Lee. “Consistent Individualized Feature Attribution for Tree Ensembles”. In: arXiv:7802.03888 [cs.LG] (Mar. 2019).

[64] 3.2.4.3.7. Sklearn.Ensemble.RandomForestClassifier — Scikit-Learn 0.23.7 Documentation. https://scikit-learn.org/stable/modules/generated/sklearn.ensemble.RandomForestClassifier.html.

[65] Python API Reference — XGBoost.XGBClassifier 7.2.0 Documentation. https://xgboost.readthedocs.io/en/latest/python/python_api.html#xgboost.XGBClassifier.feature_importances_.

[66] G. Louppe. “Understanding Random Forests: From Theory to Practice”. PhD thesis. University of Liège, Department of Electrical Engineering & Computer Science, July 2014. arXiv: 1407.7502.

[67] Shap.TreeExplainer — SHAP Latest Documentation. https://shap.readthedocs.io/en/latest/generated/shap.TreeExplainer.html.

[68] L. Breiman. “Random Forests”. In: Machine Learning 45.1 (Oct. 2001), pp. 5–32. issn: 1573-0565. doi: 10.1023/A:1010933404324.

[69] C. Molnar. “Chapter 5.5: Permutation Feature Importance”. In: Interpretable Machine Learning: A Guide for Making Black Box Models Explainable. Online book: https://christophm.github.io/interpretable-ml-book/, 2020.

[70] J. M. Spraggins et al. “High-Performance Molecular Imaging with MALDI Trapped Ion-Mobility Time-of-Flight (timsTOF) Mass Spectrometry”. In: Analytical Chemistry 91.22 (Nov. 2019), pp. 14552–14560. issn: 0003-2700. doi: 10.1021/acs.analchem.9b03612.

[71] N. H. Patterson et al. “Advanced Registration and Analysis of MALDI Imaging Mass Spectrometry Measurements through Autofluorescence Microscopy”. In: Analytical Chemistry 90.21 (Nov. 2018), pp. 12395–12403. issn: 0003-2700. doi: 10.1021/acs.analchem.8b02884.

[72] J. Lever, M. Krzywinski, and N. Altman. “Classification Evaluation”. en. In: Nature Methods 13.8 (Aug. 2016), pp. 603–604. issn: 1548-7105. doi: 10.1038/nmeth.3945.

[73] D. D. Lee and H. S. Seung. “Learning the Parts of Objects by Non-Negative Matrix Factorization”. en. In: Nature 401.6755 (Oct. 1999), pp. 788–791. issn: 1476-4687. doi: 10.1038/44565.

[74] X.-C. Xiong et al. “Feature Extraction Approach for Mass Spectrometry Imaging Data Using Non-Negative Matrix Factorization”. In: Chinese Journal of Analytical Chemistry 40.5 (May 2012), pp. 663–669. issn: 1872-2040. doi: 10.1016/S1872-2040(11)60544-6.

[75] F. Fournelle et al. “Minimizing Visceral Fat Delocalization on Tissue Sections with Porous Aluminum Oxide Slides for Imaging Mass Spectrometry”. In: Analytical Chemistry 92.7 (Apr. 2020), pp. 5158–5167. issn: 0003-2700. doi: 10.1021/acs.analchem.9b05665.

[76] M. P. Snyder et al. “The Human Body at Cellular Resolution: The NIH Human Biomolecular Atlas Program”. In: Nature 574.7777 (Oct. 2019), pp. 187–192. issn: 1476-4687. doi: 10.1038/s41586-019-1629-x.

[77] E. Ong et al. “Modelling Kidney Disease Using Ontology: Insights from the Kidney Precision Medicine Project”. en. In: 16.11 (Nov. 2020), pp. 686–696. issn: 1759-507X. doi: 10.1038/s41581-020-00335-w.

[78] R. Van de Plas et al. “Image Fusion of Mass Spectrometry and Microscopy: A Multimodality Paradigm for Molecular Tissue Mapping”. en. In: Nature Methods 12.4 (Apr. 2015), pp. 366–372. issn: 1548-7105. doi: 10.1038/nmeth.3296.

[79] M. A. Jones et al. “Discovering New Lipidomic Features Using Cell Type Specific Fluorophore Expression to Provide Spatial and Biological Specificity in a Multimodal Workflow with MALDI Imaging Mass Spectrometry”. In: Analytical Chemistry 92.10 (May 2020), pp. 7079–7086. issn: 0003-2700. doi: 10.1021/acs.analchem.0c00446.

[80] I. Jolliffe. Principal Component Analysis. Second. Statistics. Springer, 2002. isbn: 978-1-4757-1904-8.

